# A patient-driven clinicogenomic partnership through the Metastatic Prostate Cancer Project

**DOI:** 10.1101/2021.07.09.451849

**Authors:** Jett Crowdis, Sara Balch, Lauren Sterlin, Beena S. Thomas, Sabrina Y. Camp, Michael Dunphy, Elana Anastasio, Shahrayz Shah, Alyssa L. Damon, Rafael Ramos, Delia M. Sosa, Ilan K. Small, Brett Tomson, Colleen M. Nguyen, Mary McGillicuddy, Parker S. Chastain, Meng Xiao He, Alexander T.M. Cheung, Stephanie Wankowicz, Alok K. Tewari, Dewey Kim, Saud H. AlDubayan, Ayanah Dowdye, Bejamin Zola, Joel Nowak, Jan Manarite, Major Idola Henry Gunn, Bryce Olson, Eric S. Lander, Corrie A. Painter, Nikhil Wagle, Eliezer M. Van Allen

## Abstract

Molecular profiling studies have enabled numerous discoveries for metastatic prostate cancer (MPC), but they have mostly occurred in academic medical institutions focused on select patient populations. We developed the Metastatic Prostate Cancer Project (MPCproject, mpcproject.org), a patient-partnered initiative to empower MPC patients living anywhere in the U.S. and Canada to participate in molecular research and contribute directly to translational discovery. Here we present clinicogenomic results from our partnership with the first 706 MPCproject participants. We found that a patient-centered and remote research strategy enhanced engagement with patients in rural and medically underserved areas. Furthermore, patient-reported data achieved 90% consistency with abstracted health records for therapies and provided a mechanism for patient-partners to share information about their cancer experience not documented in medical records. Among the molecular profiling data from 333 patient-partners (n = 573 samples), whole exome sequencing of 63 tumor samples obtained from hospitals across the U.S. and Canada and 19 plasma cell-free DNA (cfDNA) samples from blood donated remotely recapitulated known findings in MPC and enabled longitudinal study of prostate cancer evolution. Inexpensive ultra-low coverage whole genome sequencing of 318 cfDNA samples from donated blood revealed clinically relevant genomic changes like *AR* amplification, even in the context of low tumor burden. Collectively, this study illustrates the power of a longitudinal partnership with patients to generate a more representative clinical and molecular understanding of MPC.

**Note:** To assist our patient-partners and the wider MPC community interpret the results of this study, we have included a glossary of terms in the Supplementary Materials.

## INTRODUCTION

Prostate cancer is the second most diagnosed cancer in men, with nearly 200,000 men diagnosed in 2020 alone in the U.S.^1^ Survival rates for localized disease are high, but the five-year survival rate for the over 300,000 men currently living with metastatic prostate cancer (MPC) is only 31%, representing the third leading cause of death for men^1, 2^. Because prostate cancer is largely driven by alterations to DNA, genomic sequencing studies have enabled discoveries of its molecular drivers and new therapeutic targets in both primary and metastatic clinical settings^3–6^. However, obtaining large cohorts of tumor biopsies from MPC patients for molecular study has been challenging. MPC most commonly spreads to bone, and sampling osseous lesions necessitates painful and technically challenging procedures that are not widely accessible or feasible in clinical care. Because prostate cancer can shed cell-free DNA (cfDNA) into the bloodstream, blood biopsies that sample this circulating tumor DNA have proven to be a useful alternative for the study of MPC^7, 8^.

Historically, quaternary care academic medical institutions have had the necessary infrastructure and expertise to lead clinically integrated MPC sequencing studies through clinical trials. However, the resulting clinical and genomic data is often siloed within these institutions, leading many to push for mandatory data sharing^9, 10^. These efforts, while critical to democratizing genomic research, do not directly improve access to molecular research programs and do not address underlying ethnic, socioeconomic, and geographic patient disparities in such studies, which threaten to bias findings and eventually care towards select patient populations^11–,14^. Commercial sequencing options for prostate cancer are emerging, but such approaches are often proprietary, only available to patients with appropriate insurance, and regularly inaccessible for wider research use^15–17^. Indeed, despite growing interest in clinical and research-based genomic sequencing within the MPC patient community, there are only limited mechanisms for these patients to participate in molecular profiling studies and partner with the research community to accelerate discoveries^18–20^.

We hypothesized that a patient-partnered framework that empowers MPC patients to share their biological samples, clinical histories, and lived experiences directly with researchers regardless of geographic location or hospital affiliation would lead to new clinicogenomic discoveries and begin to address demographic inequities and data access barriers in molecular studies for this disease. Thus, we established the Metastatic Prostate Cancer Project (MPCproject, mpcproject.org), a research model that leverages patient advocacy and social media to enable MPC patients to participate in genomic research remotely at no personal cost.

## RESULTS

### Development of a patient-partnered metastatic prostate cancer research model

Working with patients, loved ones, and advocates, we established an MPCproject enrollment process for men living with MPC in the U.S. and Canada (Fig. 1a). The MPCproject outreach model is community-centered and utilizes advocacy partnerships, social media campaigns, and educational initiatives to engage patients (Supplementary Fig. 1). Should they choose to register, patient-partners complete an online survey describing their experience with MPC, followed by signing electronic consent and medical release forms, which allow the MPCproject team to contact their hospitals to request medical records for abstraction and optionally archival tumor tissue for research-grade genomic sequencing (Supplementary Fig. 2). Additionally, enrolled patients can use a mailed kit to donate saliva and/or blood at routine blood draws at no cost, and these samples are sequenced to assess germline DNA and cfDNA, respectively (Supplementary Fig. 3, 4).

**Figure 1.**
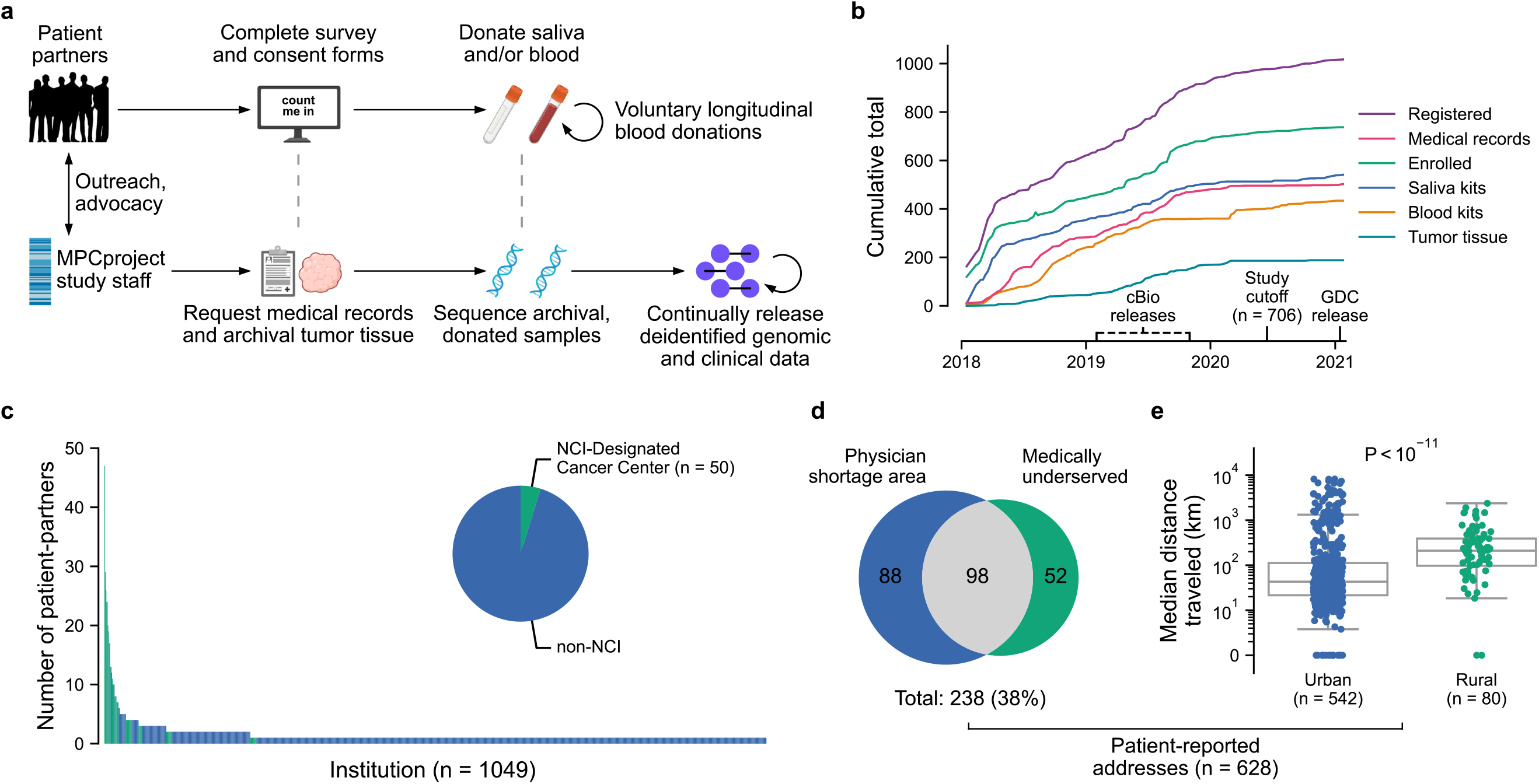
Partnering with diverse patients to enhance our understanding of metastatic prostate cancer. **a)** Summary of MPCproject enrollment process. Patients learn about the project primarily through outreach and partnered advocacy groups. If they register, patient-partners complete online intake, consent, and medical release forms, then can opt into donating saliva via a mailed kit and/or blood at routine blood draws at no charge. In parallel, MPCproject staff request medical records and archival tumor samples from patients’ medical institutions, then abstract medical information from obtained records and sequence archival tumor tissue and/or donated blood and saliva (Methods). Deidentified clinical, genomic, and patient-reported data are released on a continual, prepublication basis and deposited in public repositories. **b)** Enrollment statistics and timeline for the MPCproject. Depicted are the cumulative number of patients that began the registration process (registered), patients that completed the survey and consent forms (enrolled), patients with at least one medical record received (medical records), and blood kits, saliva kits, and archival tumor tissue received at the Broad Institute for sequencing (blood kits, saliva kits, tumor tissue, respectively). 706 patient-partners enrolled before “Study cutoff”, June 1, 2020, and are included in this study’s analyses. cBioPortal (cbioportal.org) releases include summary abstracted medical, genomic, and patient-reported data; Genomic Data Commons (GDC) releases include raw sequencing files and demographic data. **c)** Represented medical institutions among patient-partners living in the U.S. and Canada. Shown are the 1049 unique institutions (x-axis) where patient-partners report receiving care for their prostate cancer, with the number of distinct patients at each institution (y-axis). NCI-designated cancer centers are shown in green. Patient-partners that did not complete this survey question (n = 36) and institutions outside the U.S. and Canada (n = 56) are not shown. **d)** Access to medical care among patient-partners living in the U.S. Patient-reported U.S. addresses were overlapped with primary care health physician shortage areas (HPSAs) and medically underserved population/areas obtained from the Health Resources and Services Administration (HRSA.gov). Patient-partners that live in Canada (n = 30), did not provide an address (n = 40), or provided only a P.O. box (n = 8) are not shown. **e)** Patient-partners in rural areas travel farther for clinical care. Using geographic census tract information of self-reported home addresses along with USDA rural-urban continuum codes, patient-partners were categorized as living in urban or rural areas. For each patient-partner, the median Haversine round-trip distance between the zip code of their home address and that of institutions they visited was calculated (Methods). Patient-partners that live in Canada (n = 30), did not provide an address (n = 40), or provided only a P.O. box (n = 8) are not shown. *P*-value calculated via two sided Mann-Whitney U test.

Our partnership with patients is reciprocal and continuous. Patient-partners and advocates are involved in every step of the project’s design and execution—we respond directly to their feedback and keep them informed of our progress and findings (Supplementary Fig. 5). We work with men who choose to continue donating blood to help the research community understand the evolution of metastatic disease, and we regularly release prepublication, deidentified genomic and clinical data in public repositories for research use.

### Partnering with a demographically distinct patient population

To date, the MPCproject has partnered with over 1,000 patients in the U.S. and Canada and has orchestrated three public data releases (Fig. 1b). The analyses presented here are based on the 706 men from the U.S. and Canada who had enrolled (completed consent forms) as of June 1, 2020 (Supplementary Fig. 6).

Using patient-reported survey data, we assessed the geographical diversity of our patient-partners. Hailing from 49 U.S. states and 6 Canadian provinces, patient-partners reported receiving care for their prostate cancer at over 1,000 distinct medical institutions, 91% of which were reported by two or fewer patients (Fig. 1c). We found that 55% of patient-partners have never received care at an NCI-designated cancer center, where genomic research is traditionally conducted (Supplementary Table 1). These patient-partners were three times less likely to report participating in a clinical trial, indicating the understudied nature of our cohort and barriers MPC patients face in access to clinical trials (7% vs. 20%, *P* = 1 x 10^-6^, Fisher’s exact test).

Patients in rural and medically underserved areas face unique obstacles and disparities in clinical cancer care^21, 22^. To better understand the challenges faced by our patient-partners, we identified the census tracts of patient-reported U.S. home addresses and examined their geographic characteristics (n = 628/706 participants provided U.S. addresses, Methods). We found that 13% of patient-partners live in rural areas defined by the USDA, a proportion consistent with MPC patients in the U.S. generally (11%)^23^. We then examined primary care health physician shortage areas (HPSAs) and medically underserved areas (MUAs) defined by the Health Resources and Services Administration (Methods). We found that 38% of patient-partners live in HPSAs (29%) or MUAs (23%) (Fig. 1d)^24^. These proportions could not be compared with MPC patients in the U.S. due to a lack of published data, but they are significantly enriched compared to the general U.S. population (25% HPSA, 5% MUA, *P* = 0.03 and 1 x 10^-82^ respectively, Fisher’s exact test)^25, 26^. While living in a rural area was associated with being in a MUA or HPSA, 23% of MPCproject patient-partners live in urban primary care MUAs or HPSAs (*P* = 5.7 x 10^-13^, Fisher’s exact test).

We found that home addresses in rural areas were a median of 160 km farther from institutions where those patients reported receiving treatment, compared to home addresses in urban areas (*P* < 10^-11^, Mann-Whitney U test) (Methods, Fig. 1e). Although we cannot determine if home addresses changed during treatment, this suggests that patient-partners in rural areas travel significantly farther for cancer care. We did not observe significant differences in baseline clinical factors, therapies received, or likelihood to participate in a clinical trial across patients in rural areas, MUAs, or HPSAs.

The combination of the MPCproject’s online enrollment and patient-centered outreach through advocacy partnerships enabled the creation of a geographically distinct prostate cancer research program. Despite the project’s geographical diversity, however, fewer than 10% of patient-partners self-identify as non-white. While similar to existing studies, this representation remains below the proportion of minority prostate cancer patients generally (20%), a racial imbalance that has spurred new MPCproject initiatives to connect with patients of color (Supplementary Table 2, Discussion)^23^.

### Patient-reported data augment medical records to amplify patient stories

Through the patient-reported data, we sought to understand the experiences of those living with MPC. 45% of patient-partners report being diagnosed with *de novo* metastatic disease, with bone (48%) and lymph node (39%) lesions as the most common metastatic sites (Fig. 2a, b). 48% of patient-partners reported a family history of prostate or breast cancer, while 24% reported having at least one other cancer diagnosis in their lifetime, 30% of which was a non-skin form of cancer (Fig. 2c, d). The average age at diagnosis was significantly younger than the national average (61 vs. 65 years old, *P* < 10^-39^, t-test), and 24% of participants were diagnosed with early-onset prostate cancer (≤ 55 years at diagnosis, Supplementary Table 2)^27^.

**Figure 2.**
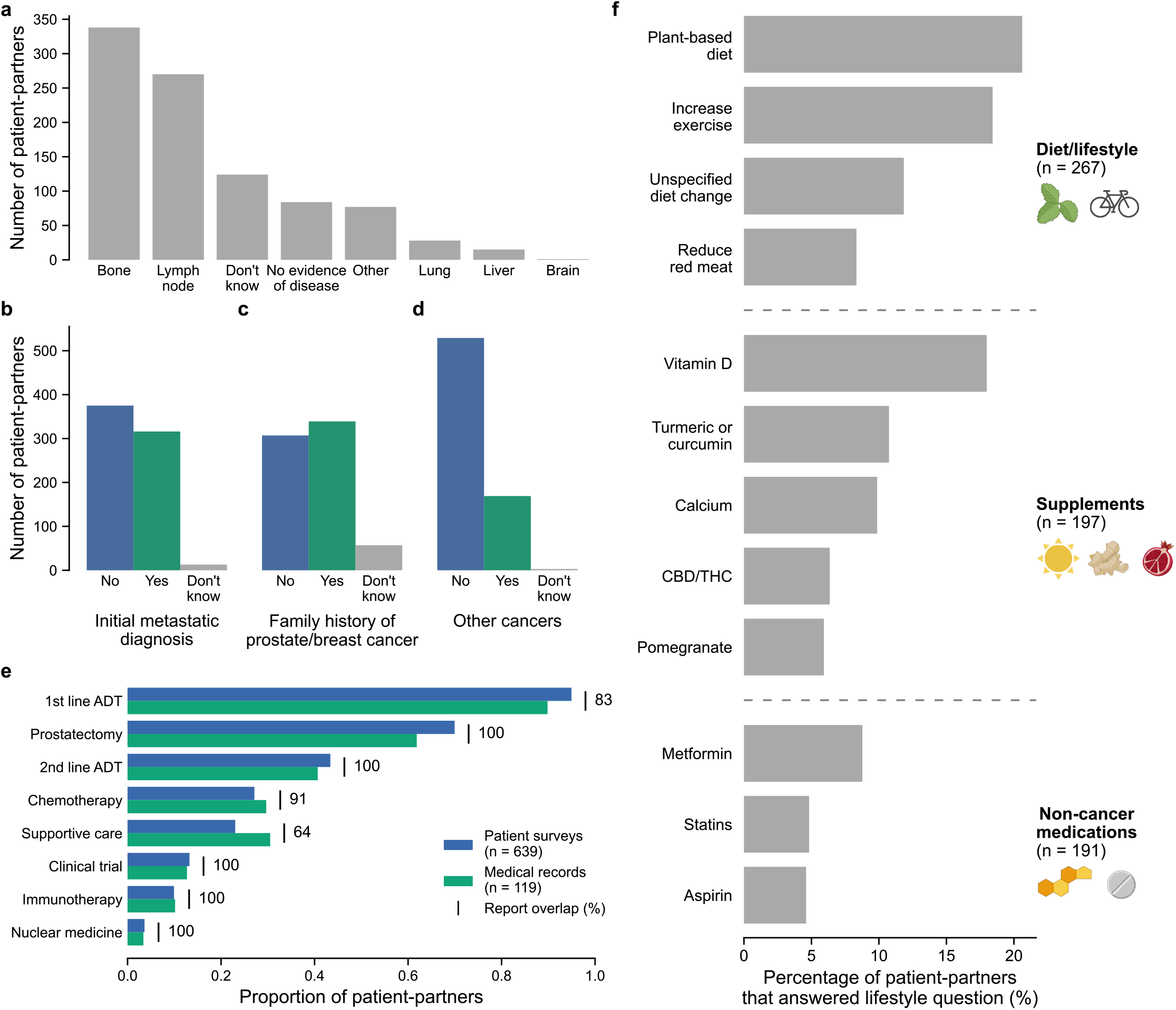
Patient voices reveal the landscape of living with metastatic prostate cancer. **a-d)** Self-reported data of 706 patient-partners related to their prostate cancer. In **a**, patient-partners were asked for the current location of their cancer. Participants were free to choose multiple if their cancer had metastasized to multiple locations. In **b-d**, responses were tabulated from questions asking patient-partners if their initial prostate cancer diagnosis was metastatic (**b**), if they have a family history of prostate/breast cancer (**c)**, or if they have ever had another cancer diagnosis (**d**). Patient-partners who did not complete these questions (n < 5) are not shown. **e)** Self-reported therapies show strong overlap with medical records. Drug categories are shown on the y-axis, with the proportion of patient-partners from each data type (patient surveys and medical records) receiving therapies of that category shown on the x-axis. In the online survey, patient-partners selected therapies they received for their metastatic prostate cancer from a list. 639/706 patient-partners reported at least one therapy and are shown. 119 of these participants also had abstracted therapy data from medical records. Report overlap refers to how often patient-partners report receiving a therapy when their medical records show that they have received that therapy, as a percentage. Only drugs available for selection in the patient survey were used in this comparison (Supplementary Table 4). **f)** Landscape of lifestyle changes for patient-partners. Participants were asked to list additional medications, alternative medications, or lifestyle changes since their diagnosis of prostate cancer. Free-text responses were manually abstracted and categorized into diet/lifestyle changes, supplements, and non-cancer medications. The y-axis shows individual instances of diet/lifestyle changes, supplements, or medications. The x-axis shows the percentage of patient-partners with that lifestyle change or taking that supplement/drug out of all patient-partners that responded to the lifestyle question (n = 456). CBD/THC: Cannabidiol/Tetrahydrocannabinol (oils, medical marijuana, etc).

We used the MPCproject’s comprehensive abstracted medical records taken from medical documentation together with patient-reported data to evaluate the treatments received in this real-world cohort (Methods, Fig. 2e). Patient-partners reported taking an average of 2.8 therapies (range 1-13) to treat their prostate cancer. 119 (17%) patient-partners had abstracted medical records at the time of writing, and there was 90% concordance between therapies noted in formal medical records and therapies reported by patients. The overlap was lowest for treatments typically given earlier in the therapeutic timeline (first line androgen deprivation therapy, 83%), supportive care therapies (64%), or treatments abandoned quickly due to side-effects (Fig. 2e). This finding illustrates the value of patient-reported data obtained via surveys for MPC, particularly in the absence of a complete medical record.

We also used the patient-reported data to assess how living with prostate cancer has changed the daily lives of our patient-partners. For example, in the survey, we asked participants to list additional medications, alternative medications, or lifestyle changes since their diagnosis of prostate cancer. 56% of patient-partners reported a lifestyle change because of living with their cancer, with the most common being a change in diet or exercise (Fig. 2f). Common nutritional supplements reported include Vitamin D and antioxidant-based supplements, while common non-cancer medications included metformin and statins. Collectively, these results demonstrate the impact of metastatic prostate cancer on patient lifestyles and that patients often pursue supplemental therapies that are not regularly documented in the medical record.

### Whole exome sequencing of a real-world MPC patient cohort

To date, we have completed molecular profiling of 573 samples from 333 patient-partners, including: ultra-low pass whole genome sequencing (ULP-WGS, average depth of 0.1x) of cfDNA from 319 donated blood samples; whole exome sequencing (WES) of cfDNA from 47 of those blood samples; WES of 106 tumor samples; and WES of 148 germline samples from donated saliva or blood buffy coat. cfDNA samples underwent WES if ULP-WGS detected a tumor fraction above 0.03 (Methods). In total, 82 exome-sequenced samples (63 tumor and 19 cfDNA) from 79 patient-partners enrolled before June 1, 2020 were included in downstream genomic analyses after assessment of sufficient tumor purity (≥10%) and coverage (Methods).

Exome sequencing from the tumor and cfDNA samples recapitulated known genomic patterns in metastatic prostate cancer (Fig. 3a). *TP53* and *SPOP* were recurrently altered, consistent with previous studies of both metastatic and primary prostate cancer (*q* < 0.1 via MutSig2CV)^3,4, 6^. In primary tumor samples from this cohort, the mutation frequency of *TP53* (30%) was more consistent with metastatic cohorts than those of primary prostate cancer^3, 6^. 17 (27%) primary tumor samples were from men diagnosed with *de novo* metastatic disease, and samples from these patient-partners were more likely to carry *TP53* mutations (*P* = 0.04, Fisher’s exact test). We also observed known patterns of copy number alteration in prostate cancer (Fig. 3a). Analysis of gene copy number alterations using GISTIC2.0 revealed recurrent amplifications of *AR* and *FOXA1*, as well as recurrent deletions of *PTEN* (*q* < 0.1)^28^. Whole-genome doubling was present in 5/63 tumor samples and 3/19 cfDNA samples, including in two tumor samples from patient-partners initially diagnosed with localized prostate cancer. In both cases, the patients were diagnosed with metastatic disease within a few months of their initial diagnosis.

**Figure 3.**
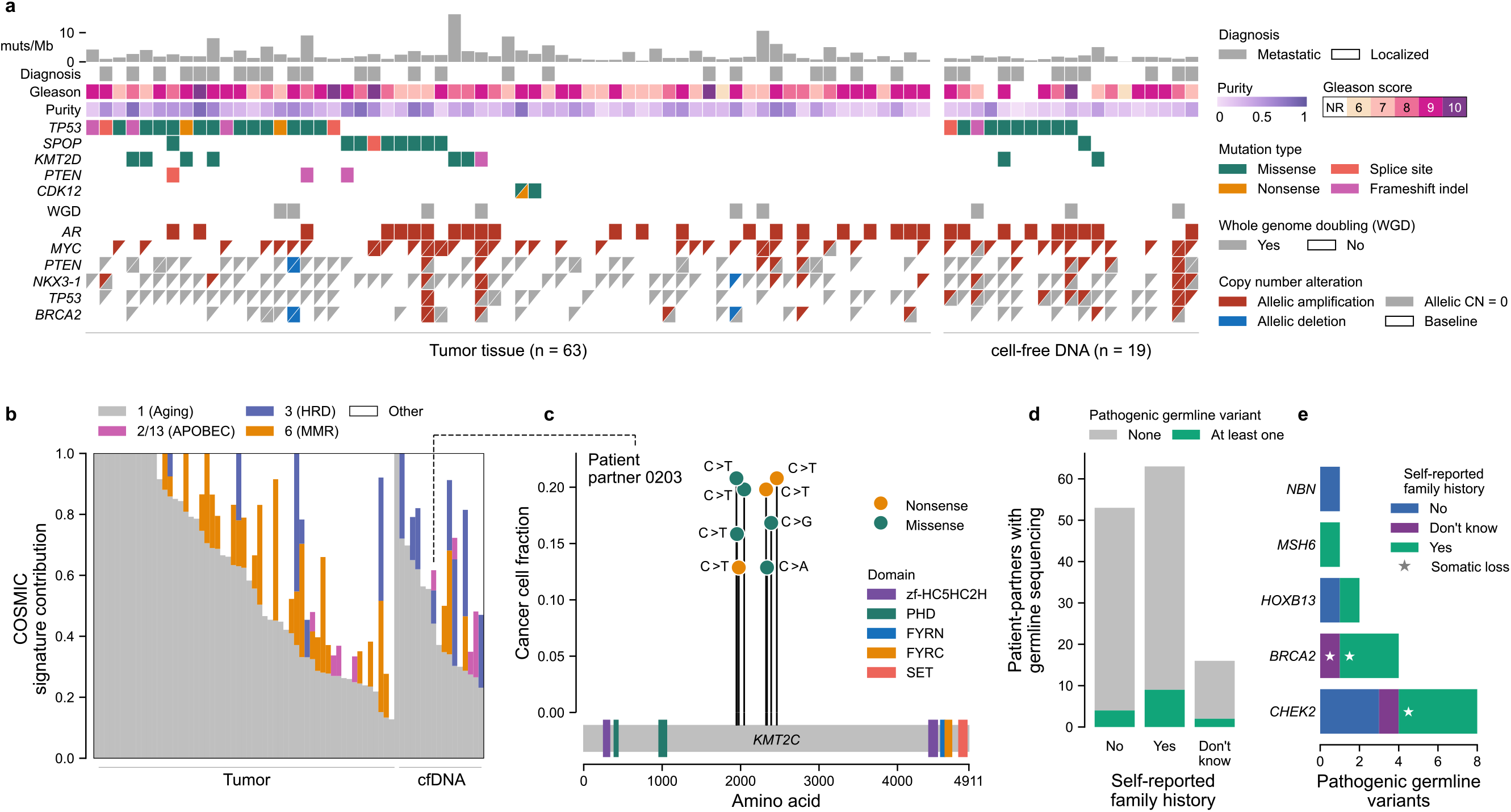
Donated tumor and cell-free DNA samples obtained through patient partnership recapitulate known genomic findings in metastatic prostate cancer. **a)** Genomic and clinical landscape of 82 sequenced samples. Columns represent samples, separated into tumor (prostate, left) and cfDNA (donated blood, right) samples, while rows represent select clinical and genomic features. Gleason scores for tumor samples are taken from the pathology report received with the sample (n = 58) or the patient-partner’s medical records (n = 5) if Gleason scores were not provided in the report. Gleason scores for cfDNA were taken from pathology reports in the medical record, with NR representing cases where a Gleason score was not reported in the medical record. Diagnosis refers to whether the initial diagnosis of prostate cancer was localized or metastatic. Multiple mutations in the same gene are represented as triangles. WGD refers to whole genome doubling. Copy number calls are allelic and defined with respect to baseline allelic ploidy (2 for samples with WGD, 1 for those without), with calls for the two alleles indicated by two triangles (except for *AR*, which has only one allele in men and so is shown as a single box). Allelic CN = 0 refers to complete allelic deletions. Allelic deletions that are not complete deletions are possible in samples with WGD. Figure created with CoMut^60^. **b)** Mutational signature analysis of sequenced samples. The relative contribution of select COSMIC v2.0 mutational signatures are shown, separated by tumor and cfDNA (donated blood) sample type^30^. APOBEC refers to signatures associated with activity of APOBEC family of cytidine deaminases (signature 2 and 13); MMR to the signature associated with deficient DNA mismatch repair (signature 6); HRD to the signature associated with homologous recombination deficiency (signature 3). Samples with too few mutations for signature analysis (< 50 mutations, n = 5 samples) are not shown. **c)** Instance of localized hypermutation (kataegis) of *KMT2C* in cfDNA from a donated blood sample. The y-axis shows the cancer cell fraction of each mutation while the x-axis shows their amino acid within *KMT2C*. Domains taken from Pfam^61^. The dotted line connects to this sample’s mutational signature profile. **d)** Germline pathogenic DNA repair alterations and their overlap with patient reported family history. Pathogenic germline alterations (as annotated by ClinVar) in genes from a select panel of DNA repair genes implicated in prostate cancer were detected in patient-partners with sequenced saliva or blood buffy coat (n = 132) (Methods; Supplementary Table 3)^62^. Survey responses to a question asking about a family history of prostate or breast cancer were tabulated and overlapped with this genomic data. Stars indicate instances where a somatic deletion also affected that gene in a tumor or cfDNA sample from that patient-partner.

To understand the mutational processes in this cohort’s exome-sequenced samples, we used a mutation-based method (deconstructSigs) to determine the contribution of COSMIC v2.0 signatures to each sample^29, 30^ (Fig. 3b, Methods). We detected the presence of aging-associated clock-like signature 1 in all samples and the presence of signature 3 (associated with homologous recombination deficiency, HRD) and signature 6 (associated with mismatch repair deficiency, MMR) in a subset of samples. These results are consistent with previous studies implicating these signatures in prostate cancer, although they likely overestimate the prevalence of signature 6 in tumor samples due to formalin-induced deamination artifacts^31, 32^. We found that the presence of signature 3 was enriched in metastasis-associated samples (cfDNA and primary tumors obtained in the metastatic setting) relative to tumor tissue from patients with strictly localized tumors at time of resection (*P <* 0.02, Fisher’s exact test). While some samples with signature 3 had alterations in *BRCA1*, *BRCA2*, or another DNA repair gene, this association was not statistically significant, potentially highlighting the presence of HRD-positive tumors without a causative molecular alteration as previously reported in studies of prostate and breast cancer^5, 33–36^.

In 10% of samples (8/82), we observed contributions from COSMIC signatures 2 and 13, which are driven by APOBEC cytidine deaminases and known to operate at a baseline level in prostate cancer^31, 37^. APOBEC-driven mutagenesis has been implicated in kataegis—rare, localized hypermutation in specific nucleotide contexts that is associated with genomic instability and increased Gleason score in prostate cancer^38, 39^. In a cfDNA sample donated by one patient-partner (patient-partner 0203), we detected eight distinct mutations within a 2 kB window in *KMT2C*, a known driver of prostate cancer (Fig. 3c)^3^. Six of these mutations were in a T(C>T)A nucleotide context, and this sample had a detectable contribution from COSMIC signature 13. We found that two pairs of the mutations, p.S1947F/p.S1954F and p.Q2325*/p.S2337Y, were each present on individual sequencing reads, confirming that these mutations existed within the same cell and strongly implicating *KMT2C* disruption through kataegis (Supplementary Fig. 7). These findings illustrate the ability to detect both frequent and rare clinically relevant molecular events in MPC across diverse contexts using a patient-partnered model.

Given the strong heritability of prostate cancer, we also sought to assess inherited germline alterations and their overlap with self-reported family history of cancer^40^. We found that among the 132 patient-partners (19%) with WES of donated saliva or blood buffy coat, 15 had pathogenic germline alterations in select genes implicated in prostate cancer heritability (Fig. 3d, Supplementary Table 3)^41^. 14% of men that reported a family history of prostate or breast cancer had at least one pathogenic germline alteration, compared to 7% of men that reported no family history, although this difference was not statistically significant (*P* = 0.38, Fisher’s exact test). The most mutated gene was *CHEK2* (8 patient-partners), followed by *BRCA2* (4 patient-partners). In three cases, we detected an accompanying somatic loss of a germline-mutated gene (Fig. 3d). These results emphasize the need to further characterize the drivers of germline susceptibility in men with MPC and to expand clinical germline testing beyond *BRCA2* in diverse clinical settings.

### Longitudinal blood biopsies enable study of tumor evolution in a patient-partnered model

Ten patient-partners had WES from both tumor tissue and cfDNA, and three patient-partners had both samples pass quality control metrics. Using the molecular data and abstracted medical records, we sought to explore the evolutionary relationships between these longitudinal samples in the context of patient clinical trajectories. Like most men with MPC, one participant, patient-partner 0495, received a diverse range of treatments between biopsy timepoints (Fig. 4a). After responding to first line anti-androgen therapy (leuprolide + bicalutamide), they took second-generation anti-androgen inhibitors (abiraterone, enzalutamide), as well as experimental radiotherapy and immunotherapy. To explore the relationship between samples, we utilized PhylogicNDT, an algorithm that clusters mutations based on their prevalence in the tumor (cancer cell fraction) into evolutionarily related subclones (Methods)^42^. In the cfDNA sample of patient-partner 0495 but not the primary tumor, we observed two distinct frameshift mutations in *ASXL2*, a gene implicated in castration-resistant metastatic prostate cancer, as well as an amplification of *AR*, a known resistance mechanism to abiraterone and enzalutamide^43, 44^. Patient-partner 0093’s tumor had clonal mutations in *TP53* and *KMT2D* but harbored an *NF2* mutation solely in the cfDNA sample. Patient-partner 0213’s tumor had a *TP53* mutation and APOBEC-associated COSMIC signature 13 detected exclusively in the cfDNA sample.

Two of these patient-partners, 0495 and 0093, were initially diagnosed with primary prostate cancer (Gleason score 4 + 3 and 5 + 4, respectively), while patient-partner 0213 was diagnosed with *de novo* metastatic disease. The primary tumor tissues of these participants were obtained at the time of diagnosis and separated from their donated blood samples by a range of years, ranging from 2 to 10 years. Despite these varied disease presentations, clinical trajectories, and biopsy timelines, we observed similar patterns of a “clonal switch” between the primary tumor and cfDNA, wherein different subclones were dominant each sample (Fig. 4b, Supplementary Fig. 8). We did not, however, observe primary tumor-specific copy number alterations, bolstering previous claims that subclonal diversification in MPC via mutations may happen after acquisition of ancestral copy number alterations (Supplementary Fig. 9)^45^. Furthermore, we observed primary tumor-specific mutations across all seven other patient-partners with both tumor and cfDNA samples, although their exact clonal structure could not be resolved due to low purity (Supplementary Fig. 10). While we cannot account for the sampling bias of tumor biopsies, these results suggest that such clonal switches may be common in the development of metastatic disease.

**Figure 4.**
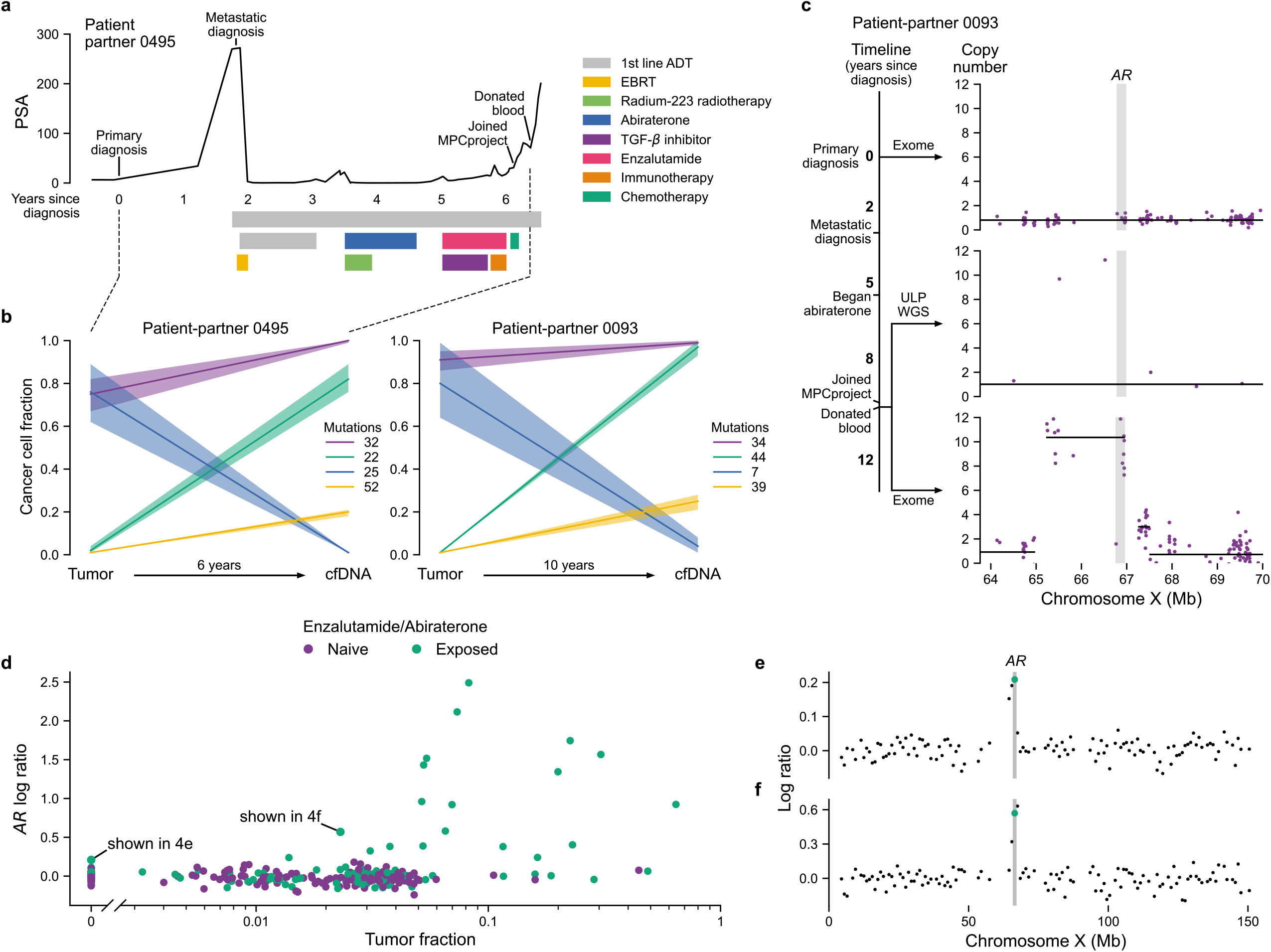
cfDNA from donated blood reveals patterns of clonal dynamics and clinically relevant genomic changes. **a)** Clinical trajectory of patient-partner 0495. This patient-partner’s prostate specific antigen (PSA) trajectory is shown on the y-axis, time in years since initial diagnosis is shown on the x-axis, and bars denote the beginning and end of therapies. EBRT—external beam radiation therapy; 1st line androgen deprivation therapy (ADT)—leuprolide and bicalutamide; immunotherapy—nivolumab; chemotherapy—cisplatin and etoposide. **b)** Tumor evolution from primary tumor to metastatic cfDNA samples. The y-axis shows the cancer cell fraction (CCF) of clonal clusters identified between tumor and cfDNA samples (x-axis). Time between samples shown on the x-axis. Colors indicate how many mutations were identified in each clone, with a 95% confidence interval around the estimated CCF. Purple represents the truncal/ancestral clone. Clusters with CCF < 0.10 across all biopsies are omitted. The clinical trajectory of patient-partner 0495 (left) is shown in **a**, while the trajectory of patient-partner 0093 (right) is shown in **c**. **c)** Emergence of *AR* amplification in patient-partner 0093 induced by anti-androgen therapy. The timeline depicts this patient’s clinical trajectory, while the plots show the absolute copy number (y-axis) of the genomic region around *AR* (x-axis, gene body shown in grey). The first plot depicts exome sequencing from the patient’s archival tumor tissue; the second and third plots depict ultra-low pass whole-genome sequencing (ULP-WGS) and exome sequencing of cfDNA from the patient’s donated blood, respectively. Individual points represent copy number of target regions (exome) or copy number of 1 Mb genomic windows (ULP-WGS). Black lines represent discrete copy number segments. **d – f)** ULP-WGS reveals clinically relevant *AR* amplifications even at low tumor fraction. Tumor fraction of 318 cfDNA samples from donated blood of 300 patient-partners with ULP-WGS sequencing is shown on the x-axis, while the log copy-ratio (logR) of the genomic interval containing *AR* is shown on the y-axis. Points are colored by whether patient-partners self-reported taking enzalutamide or abiraterone. 89 samples are shown with tumor fraction of 0 (undetectable), while 229 have nonzero tumor fractions. Two samples, one at a tumor fraction of 0 and another at a tumor fraction of 0.023, have chromosome X log copy-ratio profiles shown in **e** and **f**, respectively. The green points represent the values shown in **d,** with the genomic interval containing *AR* highlighted in grey.

In two of the three patient-partners with tumor and cfDNA samples that passed quality control, we detected the emergence of an amplification in the androgen receptor (*AR*) between the initial diagnosis and metastatic blood sample that was accurately captured using ULP-WGS of cfDNA (example patient-partner shown in Fig. 4c). This led us to examine *AR* copy number using ULP-WGS of cfDNA samples across the entire cohort, including those that did not have exome sequencing (n = 300 patient-partners, 318 samples, Fig. 4d). We found that patient-partners who reported taking enzalutamide or abiraterone had significantly higher *AR* log copy-ratios across a range of tumor fractions (*P <* 0.001, linear regression). Men who had taken enzalutamide or abiraterone also had significantly higher tumor fractions, likely reflecting a more advanced disease state and subsequent higher tumor burden in blood (*P* < 0.001, Mann-Whitney U test)^46^. We observed that *AR* amplifications are often detectable in ULP-WGS of cfDNA even when the tumor fraction is below 0.03 (Fig. 4e, f). For one patient-partner, the tumor fraction within their donated blood was inferred as undetectable, but we nevertheless observed a clear *AR* amplification (Fig. 4e). This highlights the potential efficacy of cfDNA to reveal clinically relevant changes in MPC, even in cases of very low or undetectable tumor burden. Broadly, these sequencing results illustrate the feasibility of identifying relevant genomic and evolutionary alterations from both archival tumor tissue and donated blood samples irrespective of geographical source site, enabling patient-partners to participate in genomic research at no cost and with little effort.

## DISCUSSION

Here we describe the MPCproject, a patient-driven framework for partnering with MPC patients in the U.S. and Canada to increase access to genomics research and strengthen our understanding of this disease. The online enrollment process was jointly created with patient-partners to emphasize simplicity, requiring only the completion of basic online consent and survey forms, along with optional mailed saliva and blood kits. To our knowledge, no previous effort in MPC has used patient partnership to integrated demographic, clinical, patient-reported, and genomic data from patients at a national level.

To that end, we demonstrated the feasibility of working with over 700 patient-partners, 41% of whom live in rural, medically underserved, or health physician shortage areas. We found that patient-partners living in rural areas in this study likely travel significantly farther for their cancer care, which has been shown to independently predict worse outcomes and mortality for cancer patients^47^. Furthermore, a recent study found that incomplete medical records are associated with shorter overall survival for MPC patients, particularly for those with complicated clinical histories or whose care is fragmented between institutions^48^. Our analysis of abstracted medical record data revealed a strong overlap between clinical histories represented in medical records and patient-reported data, even for patient-partners with complex treatment trajectories or who had received treatment at multiple hospitals, supporting the use of patient surveys to improve care in this disease.

We also demonstrated that tumor tissue collected from paraffin-embedded archival samples and cfDNA from donated blood samples from across the U.S. and Canada, enriched for samples not obtained from NCI cancer centers, accurately recapitulate known genomic findings in MPC, including somatic alterations, mutational signatures, germline pathogenic variants, and a rare kataegis event. There has been substantial effort in the field to identify molecular features associated with selective response to therapies like PARP inhibition and immunotherapy, including the use of mutational signatures to assess targetable HRD, MMR, and APOBEC deficiencies in cases without a causative molecular alteration^33, 49^. Our results strengthen previous findings that such signatures can be detected using cfDNA and, combined with our ability to obtain cfDNA from participants nationwide, demonstrate the scalability of a patient-partnered approach to identify and validate such genomic findings within a ‘real world’ cohort^50, 51^.

Moreover, we used archival tumor tissue and cfDNA from donated blood to reconstruct tumor phylogenetic profiles, revealing polyclonality between primary and metastatic diagnosis. Despite well-known findings of heterogeneity in both primary and metastatic prostate cancer, there is a paucity of matched primary-metastatic studies, owing mostly to the invasiveness and logistical challenges of longitudinal biopsy studies^31, 52^. Our project enables such studies paired with comprehensive clinical histories with minimal patient effort. To that end, we also found clinically relevant *AR* amplifications via low-pass WGS of cfDNA from donated blood, even at very low or undetectable tumor fractions. This result provides additional inexpensive utility to the suggested use of cfDNA tumor fraction as a clinically relevant biomarker in metastatic prostate cancer^46, 50^. We are working with patient-partners who continue to donate blood and have been able to collect multiple secondary blood biopsy kits for future longitudinal analysis.

Through feedback from patient-partners and advocates, we continue to improve the MPCproject’s design and outreach. Despite the geographic diversity of our patient-partners, we recognize that they do not reflect the racial diversity of MPC patients, a critical issue given substantial disparities in both cancer care and genomics research by race and ethnicity^11, 53, 54^. In light of structural racism and a well-founded mistrust of medical research by patients of color, this unmet disparity demands that we rethink our models of outreach and patient engagement^55^. We continue to work with community-based advocacy partners to involve communities of color, and we are building a campaign to amplify Black cancer patient voices and their lived experiences. We are also working to translate enrollment and educational materials into Spanish. In addition, a common request by our patient-partners is to enable return of clinically relevant results to participants and their physicians. While the regulatory hurdles to accomplish this are large, we recognize its importance to our patient-partners and are striving to institute return of results under this project model prospectively.

Paired with open-access clinical trials, patient-driven studies hold great promise to achieve equity and accelerate discovery in genomic research^56^. The MPCproject is part of a wider ‘Count Me In’ patient-partnered initiative (joincountmein.org) that has already yielded new findings in angiosarcoma and has expanded to metastatic breast cancer and osteosarcoma, among others^57–59^. The success of the MPCproject is based entirely on the courage and altruism of the men with whom we partner, who, in the words of one participant, hope that their “participation will help other men… and lead eventually to a cure”.

## Supporting information

Glossary of Terms

## ACKNOWLEDGEMENTS

We thank our patient-partners, caregivers, loved ones, project advisory council, and advocacy partners, without whom this project would not be possible. We would like to pay our respects to the late Jack Whelan, an MPC patient and advocate, who was instrumental in developing the MPCproject. We also thank the staff of the MPCproject, the engineering team from the Data Sciences Platform at the Broad Institute (A. Zimmer, E. Baker, S. Maiwald, P. Taheri, D. Kaplan, J. Lapan, S. Sutherland), and all members of Count Me In who work daily to ensure all MPC patients have the opportunity to participate in research. Finally, we would like to express our gratitude to the Broad Institute Cancer Program, the Broad Institute Genomics Platform, Broad Institute Communications & Development teams, and the compliance team at the Broad Institute for their support of the project. Fig. 1a and parts of Fig 2f were created with BioRender.com.

## Funding

Count Me In, Inc. Fund for Innovation in Cancer Informatics (E.M.V.), PCF-Movember Challenge Award (E.M.V.), NIH R01CA227388 (E.M.V.), U01CA233100 (E.M.V.), Mark Foundation Emerging Leader Award (E.M.V.), U.S. Department of Defense (W81XWH-21-1-0084, PC200150, SHA); Prostate Cancer Foundation (S.H.A.), Conquer Cancer Foundation of the American Society of Clinical Oncology (S.H.A.). M.X.H.: National Science Foundation (GRFP DGE1144152), National Institutes of Health (T32 GM008313).

## AUTHOR CONTRIBUTIONS

N.W., C.A.P. and E.M.V.A. conceived and designed the MPCproject with support from E.S.L. J.C., S.B., L.S., and E.M.V.A. designed and prepared the study and interpreted the data. J.C. wrote the manuscript and performed the analyses. S.B. and L.S. led study operations including tumor sample and medical record acquisition, sample sequencing, and patient coordination. L.S., B.S.T., M.D., E.A., S.S., A.L.D., R.R., D.M.S., I.K.S. oversaw medical record abstraction. S.Y.C. provided feedback on various analyses of the study and completed germline variant calling with oversight from S.H.A. S.B., L.S., J.C., B.T., M.D., M.M., and P.S.C. coordinated data releases. M.M., P.S.C., A.D., B.Z. led recent project operations. M.D. supervised early project operations. C.M.N. and E.A. led patient advocacy and outreach efforts. A.T.M.C. and S.W. oversaw early project sequencing analyses. M.X.H. provided feedback of study analyses. A.K.T. provided feedback on medical record abstractions and tissue sample collection. D.K. enabled electronic medical record searching. J.N., J.M., Major I.H.G., B.O. contributed to survey design, project development, assessment of patient criteria, and outreach strategy.

## COMPETING INTERESTS

M.X.H. has been a consultant to Amplify Medicines and Ikena Oncology. E.S.L. is currently in the process of divesting any relevant holdings. N.W. reports advisory relationships and consulting with Eli Lilly and Co.; advising and stockholding interest in Relay Therapeutics; and grant support from Puma Biotechnology. E.M.V.A. reports advisory relationships and consulting with Tango Therapeutics, Genome Medical, Invitae, Illumina, Enara Bio, Manifold Bio and Janssen; research support from Novartis and BMS; equity in Tango Therapeutics, Genome Medical, Syapse, Manifold Bio and Enara Bio; and travel reimbursement from Roche and Genentech, outside the submitted work.

## DATA AVAILABILITY

Processed, deidentified data is available on cbioportal.org (https://www.cbioportal.org/study/summary?id=prad_mpcproject_2018). Raw sequencing files are available at the Genomic Data Commons (https://portal.gdc.cancer.gov/projects/CMI-MPC). Please note that data is regularly being updated within these repositories and may not currently reflect all data generated from the project to date.

## METHODS

### Statistical computing

Except where otherwise specified, analysis and data visualization were performed with Python 3.8, SciPy v.1.5.2, Matplotlib v.3.3.2, seaborn v.0.11.0 and R v.3.5.1. All statistical tests were two-sided unless otherwise specified. The code used to generate the main figures can be found at https://github.com/vanallenlab/mpcproject-paper.

### MPCproject website

The MPCproject utilizes a website (https://mpcproject.org/) to enroll patients through an online consent and release form. The website provides information about the project and advocacy groups that have partnered with the study. The website design, messaging, and workflow were developed with direct input from patient-partners and advocates.

### Informed consent

Patients who chose to enroll in this research study are provided informed consent using a web-based consent form approved by the Dana-Farber/Harvard Cancer Center Institutional Review Board (DF/HCC Protocol 15-057B). A link to the electronic informed consent document for formal enrollment in the study (https://mpcproject.org/ConsentAndRelease.pdf) was sent to registrant emails, and upon signing, a copy of the completed form was shared. At minimum, informed consent enabled study staff to request and abstract medical records, send a saliva kit directly to patients, perform sequencing on any returned saliva samples, and release de-identified integrated clinical, genomic, and patient-reported data for research use. Patient-partners had the additional option to consent to study staff obtaining a portion of archived tumor tissue and/or a blood sample for further sequencing analysis.

### Patient-reported data

After registering, patient-partners completed a 17-question survey asking them about themselves and their disease (https://mpcproject.org/AboutYouSurvey.pdf). All questions were optional. Information on how question responses were standardized and categorized can be found in the Supplementary Methods.

### Acquisition of medical records

Medical records were obtained for patient-partners from the U.S. and Canada who completed the consent and medical release forms. Later in project development, a donated saliva or blood sample was also required. Study staff submitted medical record requests to all institutions and physician offices at which the patient reported receiving clinical care for their prostate cancer. A detailed medical record request form, along with the consent and release forms, were electronically faxed to each facility listed in a patient’s release form. Medical records were returned to the project via mail, fax, or secure online portals. If a record request was not fulfilled in six months, study staff called the hospital, and a second request was submitted, with up to three requests made. Patient-partners that communicated with study staff about changes in their treatment could request a medical record update, in which case their current hospital was again contacted for medical records. All medical records were saved in an electronic format to a secure drive at the Broad Institute.

### Acquisition of patient samples

All consented patient-partners living in the United States or Canada were mailed saliva kits with appropriate instructions, a sample tube labeled with a unique barcode, and a prepaid return box to send back the saliva sample. Samples were returned to the Broad Institute Genomics Platform, logged, and stored at room temperature (25 °C) until further sequencing.

If a consented patient-partner opted into the blood biopsy component of the study, they were sent a blood kit with instructions (https://mpcproject.org/BloodSampleInstructions.pdf, Supplementary Figure 4). Participants could take this kit to their next blood draw and request a courtesy draw by their medical provider; if a courtesy draw was not possible, patients could go to Quest Diagnostics with a complimentary voucher to have their blood drawn. Blood kits were returned free of charge to the Broad Institute Genomics Platform where they were fractionated into plasma and buffy coats and stored at −80°C. If a patient-partner did not provide a saliva sample, buffy coats were used to extract germline DNA for WES. Plasma samples continued to WES if ultra-low pass WGS detected a tumor fraction of circulating tumor DNA greater than 0.03. Some patient-partners were selected to provide additional blood samples and were sent a new consent form. If they agreed to submit another blood sample, a new blood kit was shipped.

For patient-partners that provided a germline sample and consented to the acquisition of some of their archival tumor tissue, study staff reviewed each patient’s medical records and identified available tissue (Supplementary Methods). Patient-partners were screened by the study staff to determine if they had metastatic or advanced prostate cancer based on the definition by our study. If a patient-partner had a sample that met the project’s strict requesting criteria, study staff coordinated with that hospital’s pathology department to fax a request for one H&E-stained slide as well as either 5-20 5-μm unstained slides or one formalin-fixed paraffin-embedded tissue block. Requests explicitly asked that the pathology department should not exhaust a sample to fulfill the request. Samples were sent to the MPCproject by mail. Tissue samples received as slides were labeled with unique barcode identifiers and submitted for whole exome sequencing. Tissue samples received as blocks were cut into three 30-μm scrolls per block, labeled with unique barcode identifiers, and then submitted for whole exome sequencing.

### Medical record abstraction

A data dictionary comprising 60 clinical fields with possible options was curated by trained study staff working with prostate oncologists. Electronic health records were converted to searchable PDF files using the Optical Character Recognition (OCR) engine known as Tesseract^63^. Three study staff abstractors were involved in the abstraction and QC process for each record (Supplementary Methods). If a field had lack of concordance between abstractors or there were outstanding questions, a prostate cancer oncologist reviewed the content. Whenever possible, clinical data was abstracted directly from the records. For information that’s not found, it was abstracted as ‘NOT FOUND IN RECORD’. In instances where ambiguity or incomplete data was present, inferences were made considering the whole narrative of the medical record. Incomplete dates missing the day or month are abstracted as the first day of the month or first month of the year, respectively. While all medical records will eventually be abstracted, medical records from patient-partners that received molecular sequencing of some form were prioritized for this study, resulting in 125 patient-partners with medical record abstractions, 119 of which had at least one therapy noted. In examining the overlap between patient surveys and medical record therapies, we only considered therapies that were given for metastatic prostate cancer at least one week before the patient enrolled.

### Geographic analysis

Using secure Census Bureau geocoding, we identified the census tracts of patient reported home addresses^64^. To identify patient-partners living in rural areas, this information was overlapped with rural-area continuum (RUCA) codes from the United States Department of Agriculture (USDA)^65^. Addresses with a secondary RUCA code greater than 3 were designated as rural. For comparison, the proportion of metastatic prostate cancer patients within each RUCA code from 2004 – 2017 was taken from Surveillance, Epidemiology, and End Results (SEER) using SEER*stat with the following selection table: {Site and Morphology.Site recode ICD-O-3/WHO 2008} = ‘Prostate’ AND {Stage - Summary/Historic.SEER Combined Summary Stage 2000 (2004-2017)} != ‘In situ’, ‘Localized only’, ‘Not applicable’, ‘Unknown/unstaged/unspecified/DCO’, ‘Blank(s)’^23^. To identify patient-partners living in medical shortage areas, the census tracts of home addresses were overlapped with primary care health physician shortage areas (HPSA) and medically underserved areas (MUA) defined by the Health Resources and Services Administration (HRSA)^25^. Addresses were labelled as existing within a MUA if they were designated as within a medically underserved area or population and as existing within a HPSA if they were designated as within a primary care HPSA. Published geographic datasets of cancer patients (e.g., SEER, NPCR) do not contain census-tract resolved data or summary results of MUA/HPSA status, so for comparison we instead used the total U.S. population living in HPSAs and MUAs, taken from HRSA, divided by the entire U.S. population taken from the U.S. Census^25, 26^. To calculate appointment distances, we calculated the round-trip Haversine distances between the zip code of home addresses and the zip code of reported institutions.

### Whole exome sequencing analysis

Whole exome sequences were captured using Illumina technology and the sequence data processing and analysis was performed using Picard and FireCloud pipelines on Terra (https://terra.bio/) (Supplementary Methods). The Picard pipeline (http://picard.sourceforge.net) was used to produce a BAM file with aligned reads. This includes alignment to the GRCh37 human reference sequence using BWA^66^ and estimation and recalibration of base quality score with the Genome Analysis Toolkit (GATK)^67^. Somatic alterations for tumor samples were called using a customized version of the Getz Lab CGA WES Characterization pipeline (https://portal.firecloud.org/#methods/getzlab/CGA_WES_Characterization_Pipeline_v0.1_Dec2 018/) developed at the Broad Institute. Briefly, MuTect v1.1.6 algorithm was used to identify somatic mutations^68^. Somatic mutation calls were filtered using a panel of normals (PoN), oxoG filter and an FFPE filter to remove artifacts introduced during the sequencing or formalin fixation process^69^. Small somatic insertions and deletions were detected using the Strelka algorithm^70^. Somatic mutations were annotated using Oncotator^71^. Recurrently altered mutations were identified using MutSig2CV^72^. To define somatic copy ratio profiles, we used GATK CNV^67^. To generate allele-specific copy number profiles and assess tumor purity and ploidy, we used ABSOLUTE and FACETS^73, 74^. Final segmentation calls were taken from ABSOLUTE, except for the X chromosome, which was taken from FACETS. We utilized GISTIC2.0 to identify significantly recurrent amplification and deletion peaks^28^. For determining allele-specific copy number alterations, we assessed the absolute allelic copy numbers of the segment containing each gene. Mutation burden was calculated as the total number of mutations (non-synonymous + synonymous) detected for a given sample divided by the length of the total genomic target region captured with appropriate coverage from whole exome sequencing.

### Whole exome sequencing quality control

Samples with average coverage below 55x in the tumor sample or below 30x in the normal sample were excluded. Samples with purity < 0.10 from both ABSOLUTE and FACETS were excluded. DeTiN was applied to samples to estimate the amount of tumor contamination in the normal samples; samples with TiN (tumor in normal) > 0.25 were excluded^75^. ContEst was applied to measure the amount of cross-sample contamination in samples; samples with contamination > 0.04 were excluded^76^. The Picard task CrossCheckFingerprints was applied to determine sample mixups; samples with Fingerprints LOD value < 0 were excluded^77^. Samples which passed quality control were submitted to cBioPortal and GDC.

### Ultra-low pass whole genome sequencing analysis

ichorCNA was used to assess the tumor fraction in cfDNA samples that completed ultra-low pass whole genome sequencing^50^. The log copy ratio of *AR* was assessed by the log copy ratio of the genomic interval containing *AR.* This value could not consistently be converted to absolute copy number due to the low tumor fractions of many samples.

### Mutational signature analysis and kataegis

Mutational processes in our cohort were determined using deconstructSigs with default parameters applying COSMIC v2 signatures as the reference with a maximum number of signatures of 6^29, 30^. A signature was assessed as present if the signature contribution was greater than 6%. Because tumor samples were formalin-fixed and paraffin embedded (FFPE), a process known to introduce stranded mutational artifacts in specific nucleotide contexts, we used a filter to remove likely FFPE artifacts according to nucleotide context and strand bias before using deconstructSigs^78^. We also tried to assess the colocalization of the kataegis event with structural variant breakpoints but were limited by targeted sequencing in exomes and low coverage in ULP-WGS. *KMT2C* and its surrounding region were not copy number altered in the sample with kataegis. Kataegis was not identified in any other sample.

### Association of DNA-repair alterations and presence of signature 3

Alterations in a select list of genes previously implicated in DNA-repair in prostate cancer were examined (Supplementary Table 3). An alteration was considered if there was a somatic single-copy deletion, double deletion, nonsense mutation, missense mutation, frameshift indel, or splice site mutation. An alteration was also considered if there was a pathogenic germline alteration, denoted by “Pathogenic” in ClinVar^62^.

### Germline variant discovery

To call short germline single-nucleotide polymorphisms, insertions, and deletions from germline WES data, we used DeepVariant (v0.8.0)^79, 80^. Specifically, we used the publicly-released WES model (https://console.cloud.google.com/storage/browser/deepvariant/models/DeepVariant/0.8.0/Deep Variant-inception_v3-0.8.0+data-wes_standard/) to generate single-sample germline variant call files using the human genome reference GRCh37(b37). We filtered variants with bcftools v1.9 to only keep high-quality variants annotated as “PASS” in the “FILTER” column. The high-quality variants were merged into single-sample Variant Call Format (VCF) files using CombineVariants from GATK 3.7 (https://github.com/broadinstitute/gatk/releases). To decompose multiallelic variants and normalize variants, we used the computational package vt v3.13 (https://github.com/atks/vt). Lastly, germline variants were annotated using the VEP v92 with the publicly-released GRCh37 cache file (https://github.com/Ensembl/ensembl-vep)81. Germline variants were denoted as pathogenic if they appeared as “Pathogenic” in ClinVar (Dec 2019 version)^62^.

### Phylogenetic analysis

To compare mutations between distinct samples (tumor and cfDNA) from the same patient, we used a previously described method designed to recover evidence for mutations called in one sample in all other samples derived from the same individual^82^. In brief, the ‘force-calling’ method uses the strong prior of the mutation being present in at least one sample in the patient to more sensitively detect and recover mutations that might otherwise be missed. A mutation was deemed tumor/cfDNA specific if there were no force-called reads that supported the mutation in the other sample, although this process underestimates the proportion of shared mutations in low purity tumors. The cancer cell fraction (CCF) of mutations were defined using ABSOLUTE, which calculates the CCF based on variant allele frequency, purity, and local allelic copy number^73^. To reconstruct tumor phylogenies, we used PhylogicNDT, which clusters mutations into subclones across multiple samples based on their underlying similar CCFs^42^.

### Data releases

The MPCproject releases de-identified clinical, patient-reported and research-grade genomic data into public repositories, such as cBioPortal (https://www.cbioportal.org/study/summary?id=prad_mpcproject_2018) and the Genomic Data Commons (https://portal.gdc.cancer.gov/projects/CMI-MPC), at regular intervals and pre-publication. Data is processed and formatted as required by each repository’s guidelines. All patient identifiers are stripped prior to data deposition to protect patient privacy. On the MPCproject data release webpage (https://mpcproject.org/data-release), patients can access project data, additional information about the data, list of common terms used in research, methods used to generate the data, and an email address for any additional data-related questions.

