## Supplementary material for "A patient-driven clinicogenomic partnership through the Metastatic Prostate Cancer Project": Glossary of Terms

### MPCproject glossary

**Note:** This glossary is intended to assist our patient-partners and the wider MPC community understand the results of this study. The definitions are therefore optimized for readability.

#### Clinical terms

- **Metastasis** – When a cancer spreads outside the area of the body where it started. In prostate cancer, metastasis occurs when the cancer spreads outside the prostate gland.
- **Primary site** – The area in the body where the cancer started. In metastatic prostate cancer, the prostate is the primary site. A cancer that has not spread is sometimes called primary cancer.
- **De novo metastasis** – Cancer that is already metastatic when it is diagnosed.
- **Treatment resistance** – When a cancer continues to spread during treatment, even if it responded initially.
- **Molecular profiling study** – Research that studies the properties of tumor samples.
- **Translational research** – Research that is focused on applying research to clinical settings and medicine.
- **Genomic data** – Data that is generated by sequencing (looking at) DNA. "Genomic" is a term that generally refers to the study of DNA.
- **Clinicogenomic** – Combining clinical data with genomic data.
- **Abstraction (or abstracted medical records)** – The act of reading medical records and summarizing the information of interest.
- **Patient-reported data** – Data collected from surveys completed by patients.

#### Genomic terms

- **DNA** – The genetic material within cells that instructs the cells how to operate. Every individual has his or her own unique DNA.

- **Mutation** – A change in the DNA sequence. This in turn can change the function of the DNA and may disrupt how cells normally operate.

- **Gene** – A piece of DNA that has a specific function. Cancer usually arises because cells acquire mutations that disrupt the function of genes, leading to errors such as uncontrolled growth in the form of a tumor. For each gene, there are two copies (alleles), one inherited from each parent.

- **Protein** – The functional form of a gene. A gene's DNA is transformed ("expressed") into a protein by the cell to perform its specific function. Mutations in genes can result in dysfunctional proteins, which disrupt the cell's natural processes.

- **Somatic mutation** – A mutation that happens during a person's lifespan. Somatic mutations are not obtained from parents and are not passed to offspring. In cancer, somatic mutations are gained only by cancer cells. Somatic mutations are typically detected by sequencing a biopsy of the cancer tissue.

- **Germline mutation** – A mutation that can be passed from parents to offspring. People are born with germline mutations, which are present in all cells of their body. Some germline mutations increase the risk of getting cancer, although having one does not guarantee a person will get cancer. Germline mutations may result in a family history of cancer, like a germline mutation in the gene *BRCA2* in prostate cancer. They are detected by sequencing a blood or saliva sample.

- **Copy number alteration –** Changes in the number of copies of a gene above or below the normal 2 copies. This can be either an amplification or a deletion (see below).

- **Gene amplification** – An increase in the number of copies of a gene above two. This causes a gene to be expressed more than usual and is a common way cancer cells become resistant to drugs. In prostate cancer, for example, the androgen receptor gene (*AR*) is often amplified, which leads to therapy resistance against drugs like enzalutamide (Xtandi) and abiraterone (Zytiga).

- **Gene deletion** – The loss of all or a part of a gene, reducing, eliminating, or otherwise changing the gene's function. Genes that prevent cancer growth, such as *TP53*, are often deleted in cancer.

- **Copy ratio –** A metric that quantifies changes in the number of copies of a gene. It is based on how much DNA of that gene is present in a sample.

- **Mutational signatures** – Somatic mutations can follow patterns, known as mutational signatures, that suggest a common process that caused them. Some patterns are normal and associated with aging (signature 1), but others are associated with a mutation repair process in the cell that has gone wrong (signature 3 and 6, see HRD/MMR).

- **HRD (Homologous recombination deficiency)** – Cells normally repair breaks in DNA through a process called homologous recombination. Mutations in genes involved in DNA repair can hinder this process, leading to HRD. For example, in prostate cancer, germline or somatic mutations in the gene *BRCA2* can lead to HRD, and the drug Olaparib (Lynparza) is approved to treat these *BRCA2* mutated prostate cancers. Mutational signature 3 has been associated with HRD.

- **MMR (**mismatch repair deficiency) – When cells divide, they replicate their DNA, and sometimes errors can occur during this process (mutations called "mismatches"). Usually, there are genes that help the cell repair these errors. If these repair genes are disrupted, a cell's DNA can be highly prone to mutations, leading to MMR. Mutational signature 6 has been associated with MMR.

- **APOBEC –** Stands for "apolipoprotein B mRNA editing enzyme, catalytic polypeptide–like". APOBEC proteins are a family of proteins that normally help the body fight viral infections. In cancer, they can become dysregulated and lead to increased mutations in the cancer cell's DNA.

- **Kataegis –** An event in which many mutations occur close together in a highly patterned form.

- **Whole genome doubling –** Normal cells have 2 copies of each gene. Errors can occur during a cell's life cycle where all the DNA is doubled, leaving the cell with 4 copies of each gene. Whole genome doubling is associated with advanced prostate cancer.

- **Cancer evolution (phylogenetics) –** Cancer cells follow the same evolutionary principle of natural selection ("survival of the fittest") that can be applied to animals. Cancer cells that acquire mutations that help them survive will replicate more than those that do not. Scientists can map these mutations over time to study how a cancer changes (evolution).

- **Clonality (clonal, subclonal, polyclonal) –** Cancer cells descend from a common ancestor cell but acquire mutations as they multiply. Because DNA is passed from a parent cell to its two offspring, all cancer cells will share mutations that were acquired early in that cancer's development. Such mutations are called clonal mutations. Mutations that are acquired later by only a subset of cancer cells are called subclonal. A polyclonal tumor has many subclonal mutations.

Sequencing terms

- **Genomic sequencing** – A process that allows researchers to "read" DNA extracted from a sample.

- **Types of sequencing (WES, ULP-WGS)** – Whole exome sequencing (WES) reads only the DNA that is contained within genes (the "exome", about 1% of all DNA). Whole genome sequencing (WGS) reads all the DNA (the "genome"). Ultra-low pass WGS (ULP-WGS) reads all the DNA at a shallow level, enough to assess tumor fraction in cell-free DNA.

- **Archival tumor sample –** A piece of tissue taken from the prostate, usually at the time of diagnosis or via a prostatectomy. These samples are preserved in a substance called paraffin and stored in hospitals. In this project, archival tumor samples are requested from hospitals, removed from paraffin, and then sequenced to read the tumor's DNA.

- **Purity** – When a biopsy is taken, not all of the tissue is cancerous. Purity refers to the percentage of a biopsy that is cancerous. Low purity tumors are more difficult to study, as they contain fewer cancer cells. In the context of cell-free DNA/liquid biopsies, purity is also called tumor fraction.

- **Formalin induced deamination artifact –** Archival tumor samples are stored in paraffin using a chemical called formalin. When in formalin, DNA can gain mutations that were not originally in the cancer. These are called "artifacts" and will be detected in sequencing but are not relevant for the cancer. We use a filter to remove these artifacts from sequencing data.

- **Cell-free DNA (cfDNA) –** As cells die, they can shed DNA into the bloodstream, resulting in cell-free DNA. Metastatic prostate cancer cells can sometimes shed enough DNA to be detected by sequencing (circulating tumor DNA). As a result, cfDNA is increasingly being used to monitor and evaluate metastatic prostate cancer.

- **Liquid biopsy** – A blood draw designed to assess cell-free tumor DNA. In this project, these biopsies are received by participants via FedEx (free of charge) and then sequenced to detect circulating tumor DNA.

- **Tumor fraction –** The percentage of cancer DNA compared to normal DNA in a cell-free DNA sample (0% - 100%).
