## Supplementary Figures and Tables for "A patient-driven clinicogenomic partnership through the Metastatic Prostate Cancer Project"

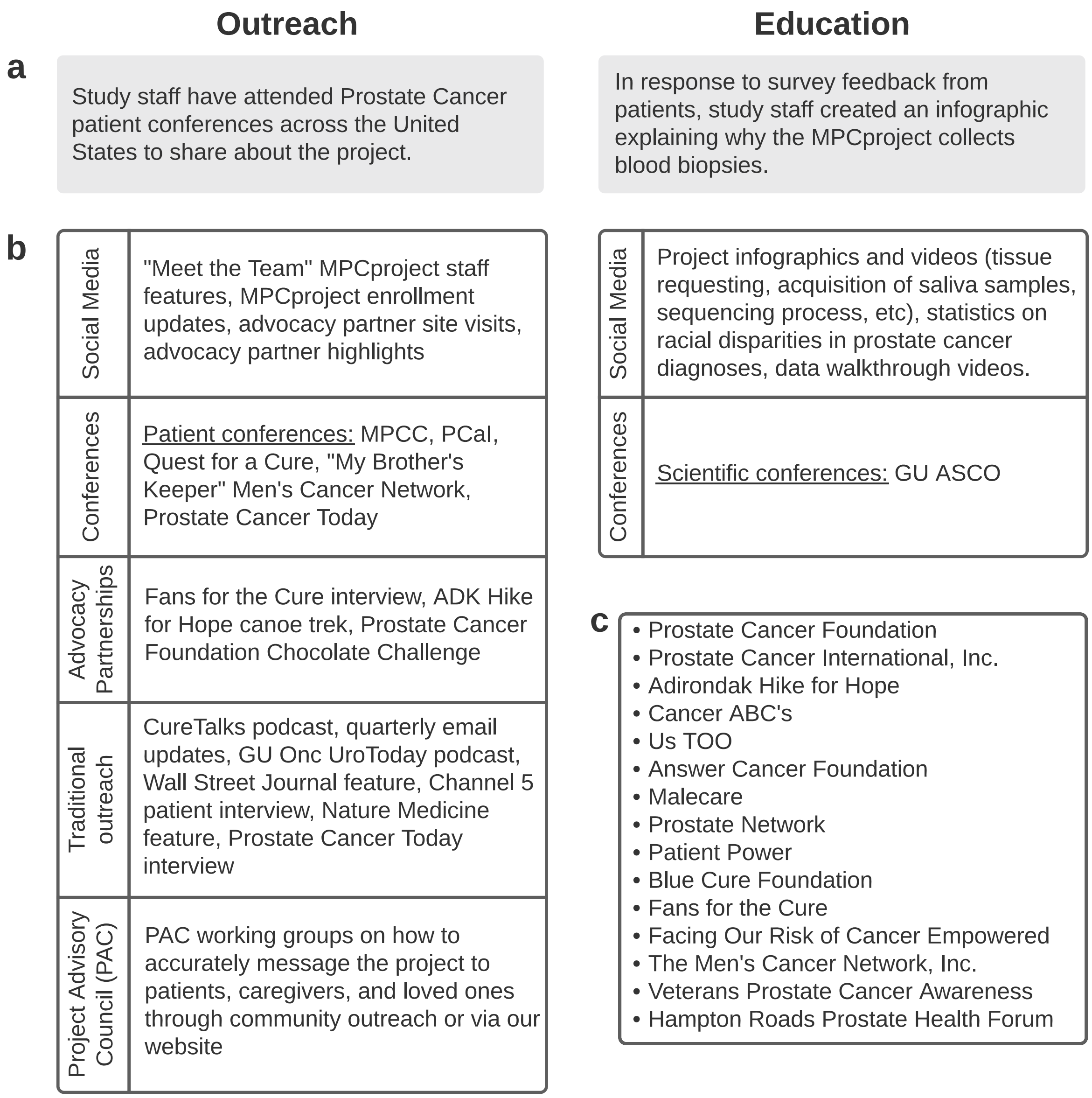


**Supplementary Figure 1. MPCproject education and outreach initiatives reach patient-partners across the country.**

**a)** Education and outreach spotlights. Study staff attend and present at patient conferences to share information about the MPCproject with the extended prostate cancer community. Conference tables have example sample kits, brochures, and a mailing list sign-up to learn more. For patients who follow the MPCproject on social media, study staff create online polls to identify educational content important to the community. One such poll revealed interest in learning about the biological significance of liquid biopsies and why the project collects them. **b)** Select examples of outreach and education initiatives. As a result of the decentralized, online nature of the study, the MPCproject uses diverse modes of education and outreach to reach patient-partners. **c)** The MPCproject partners with patient advocacy groups across the United States and Canada. Advocacy partners help encourage patient participation in the project as well provide ongoing input regarding the design and implementation of the project overall.

**a**

Please fill out as much as you can. All questions are optional. You can return at any time with the link sent to you by email.

1. When were you first diagnosed with prostate cancer? If you do not remember the month, you can enter just the year.

   Choose month... | Choose year...

2. When you were first diagnosed, were you diagnosed with advanced or metastatic prostate cancer (prostate cancer that has spread beyond the prostate, including biochemical recurrence)?

   ◉ Yes
   ○ No
   ○ I don't know

3. Did you receive local treatment to your prostate when you were first diagnosed (local treatment includes surgery, radiation, or cryotherapy)?

   ◉ Yes
   ○ No
   ○ I don't know

4. Have you had your entire prostate surgically removed (known as a prostatectomy)?

   ◉ Yes
   ○ No
   ○ I don't know

5. Where is your prostate cancer currently located (check all that apply)?

   ☑ Lymph Node
   ☐ Bone
   ☐ Liver
   ☐ Lung
   ☐ Brain
   ☑ Other

   Please provide details

   ☐ No Evidence of Disease (NED)
   ☐ I don't know

6. For your advanced prostate cancer (prostate cancer that is outside of the prostate), please check off all therapies that you have previously received or are currently receiving (Check all that apply)

   Hormones
   ...
   Chemotherapy
   ...
   Other Therapy
   ...

   See Supplementary Table 4 for therapy list

   Experimental/Clinical Trial

   ☑ Experiment/Clinical Trial

   Please provide details

   ☑ Other

   Please provide details

7. Please list additional medications, alternative medications, you've taken or lifestyle changes that you've made since your diagnosis with prostate cancer.

8. Have you had any other types of cancer?

   ◉ Yes
   ○ No
   ○ I don't know

9. What other cancer(s) have you had?

10. Do you have any family history of prostate and/or breast cancer?

    ◉ Yes
    ○ No
    ○ I don't know

11. How did you find out about this project?

12. Is there anything else you would like us to know about your prostate cancer?

13. Do you consider yourself Hispanic, Latino or Spanish?

    ◉ Yes
    ○ No
    ○ I don't know

14. What is your race (select all that apply)?

    ☐ American Indian or Native American
    ☐ Japanese
    ☐ Chinese
    ☐ Other East Asian
    ☐ South East Asian or Indian
    ☐ Black or African American
    ☐ Native Hawaiian or other Pacific Islander
    ☐ White
    ☐ I prefer not to answer
    ☑ Other

    Please provide details

15. In what year were you born?

    Choose year...

16. What country do you live in?

    Choose country...

17. What is your ZIP or postal code?

    Zip Code

I understand that the information I entered here will be stored in a secure database and may be used to match me to one or more research studies conducted by the Metastatic Prostate Cancer Project. If the information that I entered matches a study being conducted by the Metastatic Prostate Cancer Project, either now or in the future, I agree to be contacted about possibly participating. I understand that if I would like my information deleted from the database, now or in the future, I can and my information will be removed from the database.

SUBMIT

**Supplementary Figure 2. MPCproject About You Intake Survey**

**a)** After registering, patient-partners complete an online intake survey detailing their experience with metastatic prostate cancer (https://mpcproject.org/AboutYouSurvey.pdf). All questions are optional. Questions were developed in collaboration with patient-partners and practicing prostate cancer oncologists. For a full list of therapies for question 6, see Supplementary Table 4. The survey responses above are shown as an example and do not represent any specific patient-partner's responses.

a b

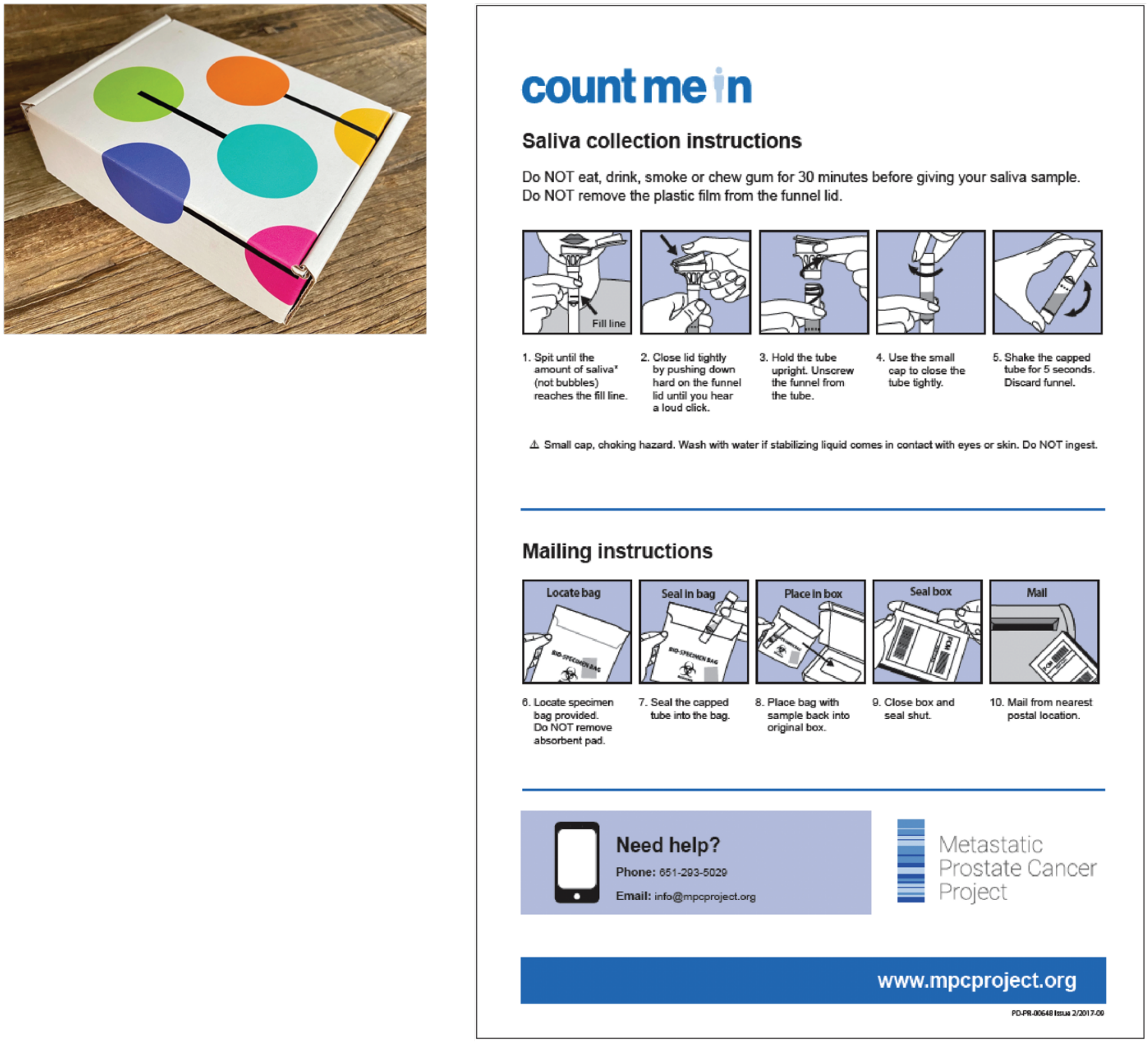


**Supplementary Figure 3. MPCproject remote saliva donation kit**

**a)** Enrolled patients in the U.S. and Canada are mailed a saliva kit. Each kit comes with a tube for saliva donation and a prepaid FedEx return envelope. All components of the kit, including the box itself, contain a unique, nonidentifiable barcode associated with the patient-partner. Acting on feedback about privacy from patient-partners and advocates, boxes are kept nondescript to avoid identifying the recipient as a patient with prostate cancer.

**b)** Saliva kit instructions. These instructions are included in the box itself, and patient-partners can contact the MPCproject study team for additional assistance if necessary.

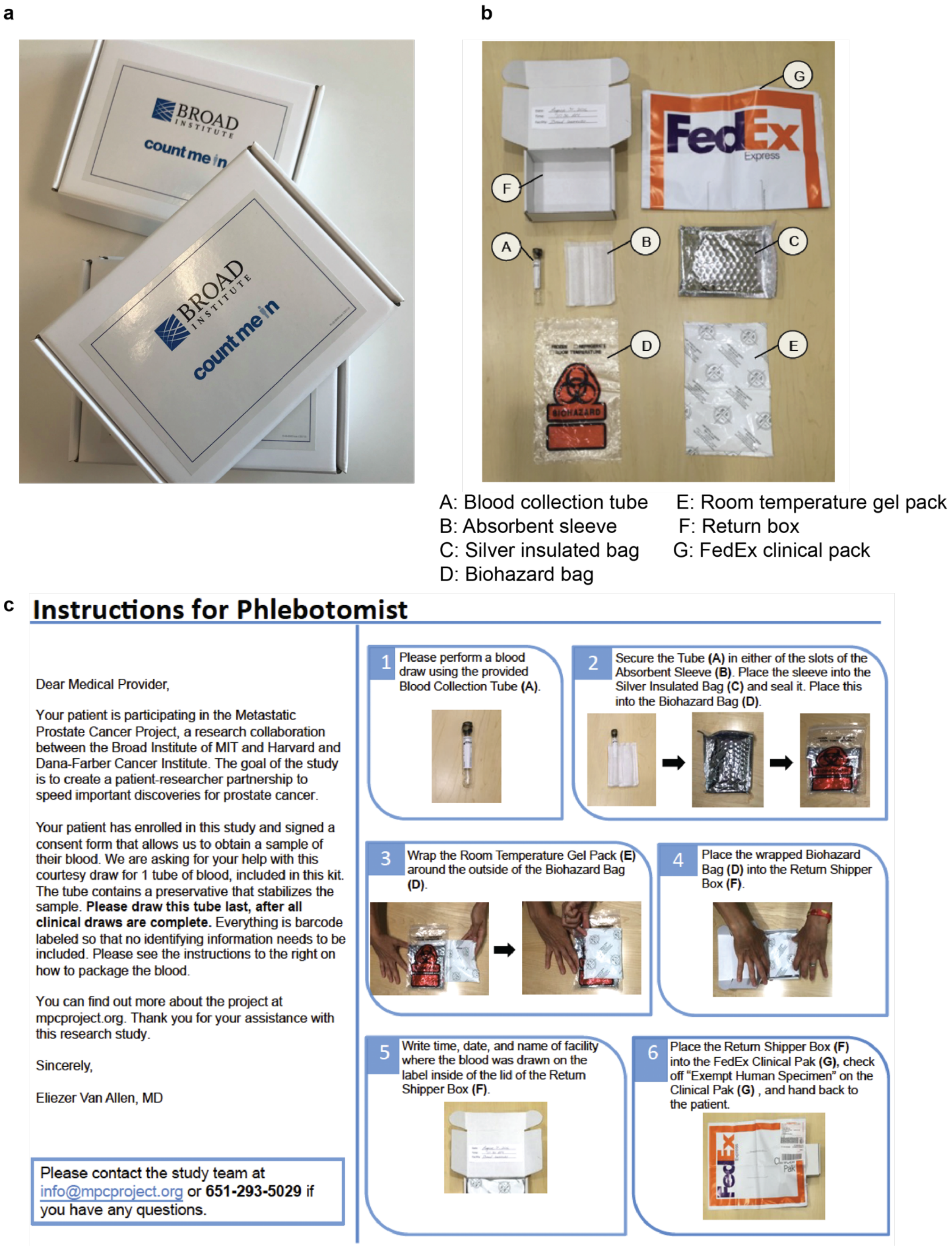


**Supplementary Figure 4. MPCproject blood donation kit**

**a)** If they consented to donate blood on their online survey, patient-partners are mailed a blood kit. Each kit comes with a tube for blood donation, instructions for use, and a unique, nonidentifiable barcode. Acting on feedback about privacy from patient-partners and advocates, boxes are kept nondescript to avoid identifying the recipient as a patient with prostate cancer.
**b)** Composition of blood donation kit. This graphic is included within the blood donation kit.
**c)** Instruction for healthcare providers. Patient-partners provide these instructions to their healthcare provider or phlebotomist at regular, standard of care blood draws. A courtesy draw is requested, free of charge, but if this is not available, patient-partners can also visit a local Quest Diagnostics lab with a free voucher for a blood draw. After completion, the kit is placed within the prepaid FedEx envelope and mailed to the Broad Institute where it is kept for sequencing.

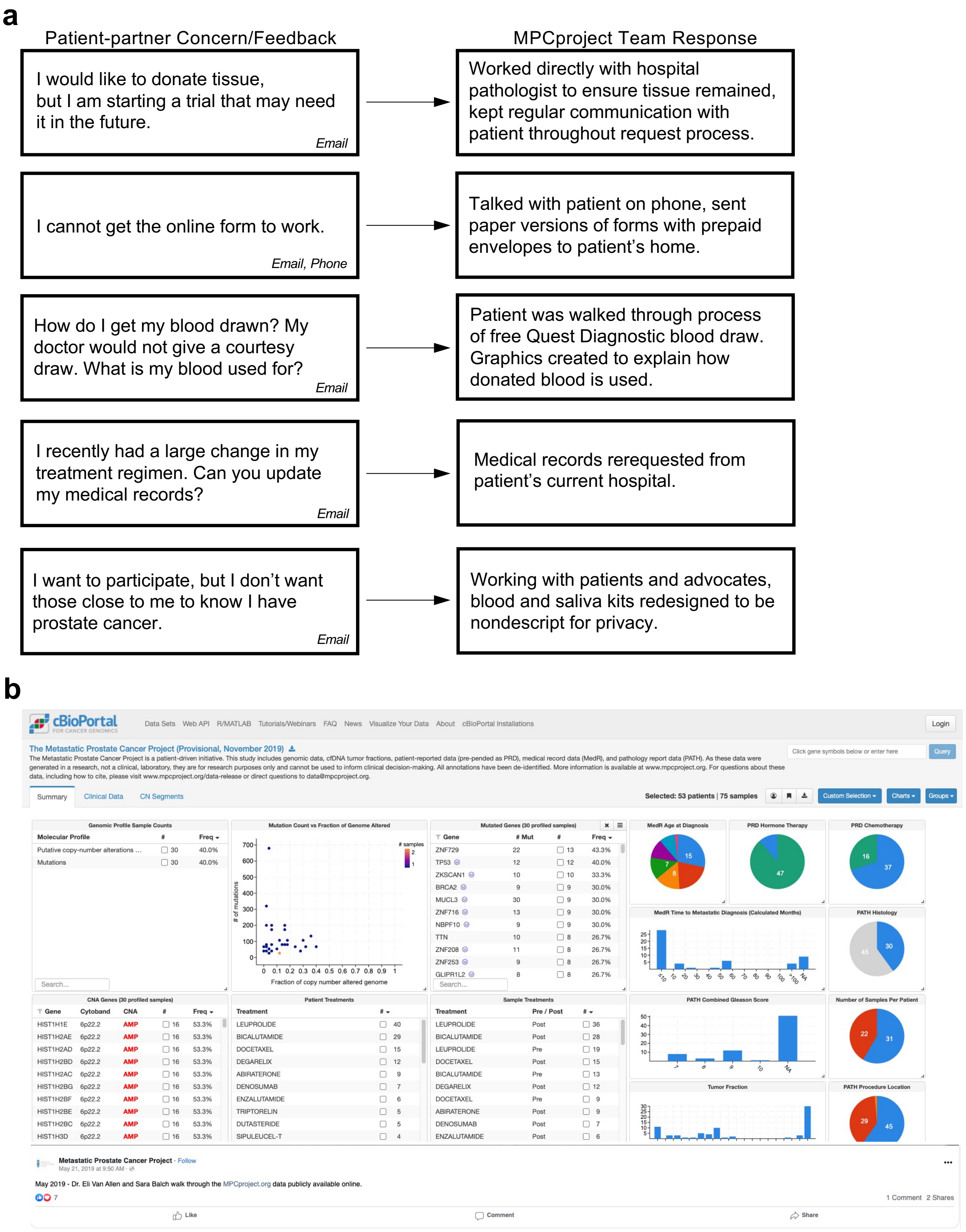
a
Patient-partner Concern/Feedback
MPCproject Team Response
I would like to donate tissue, but I am starting a trial that may need it in the future.
Email
Worked directly with hospital pathologist to ensure tissue remained, kept regular communication with patient throughout request process.
I cannot get the online form to work.
Email, Phone
Talked with patient on phone, sent paper versions of forms with prepaid envelopes to patient's home.
How do I get my blood drawn? My doctor would not give a courtesy draw. What is my blood used for?
Email
Patient was walked through process of free Quest Diagnostic blood draw. Graphics created to explain how donated blood is used.
I recently had a large change in my treatment regimen. Can you update my medical records?
Email
Medical records rerequested from patient's current hospital.
I want to participate, but I don't want those close to me to know I have prostate cancer.
Email
Working with patients and advocates, blood and saliva kits redesigned to be nondescript for privacy.
b
cBioPortal
The Metastatic Prostate Cancer Project (Provisional, November 2019)
Summary
Clinical Data
CN Segments
Selected: 53 patients | 75 samples
Metastatic Prostate Cancer Project
May 2019 - Dr. Eli Van Allen and Sara Balch walk through the MPCproject.org data publicly available online.

**c**

Metastatic Prostate Cancer Project

Dear MPCproject Mailing List,

We are writing to share another quarterly update with the patients, caregivers, scientists, and advocacy partners engaged in the Metastatic Prostate Cancer Project (MPCproject.org). We are tremendously grateful for your continued support. Below you will find some recent news from the MPCproject.

**The Numbers**

Over 745 patients have clicked "Count Me In" to register for the project. Thanks to your generosity, we have received:

- 662 medical records
- 417 saliva kits
- 329 blood samples

**Understanding Our Data**

The MPCproject recently released a video of Dr. Eli Van Allen walking through the genomic and clinical data we have released thus far on cBioPortal. We will continue to release data as it is generated, pre-publication. If you have any questions regarding the data currently available, please email us at.

**Supplementary Figure 5. Working directly with patients in the MPCproject**

**a)** Examples of feedback from patient-partners and the response of the project team. In each case, patient-partners contacted the MPCproject office with concerns, questions, or feedback. The MPCproject study staff maintains regular contact with patient-partners that have questions and creates infographics and educational materials based on common questions.

**b)** Walkthrough of initial MPCproject data on cBioPortal. When the project's first data release happened on cBioPortal, Dr. Van Allen and the study team recorded a walkthrough (https://m.facebook.com/watch/?v=471939353546532) explaining the shared MPCproject data to patient-partners.

**c)** Quarterly email updates. An example of a quarterly update sent four times a year to patient-partners, loved ones, and advocates on the MPCproject mailing list. These emails explain study progress, how to interpret data releases, and new project initiatives.

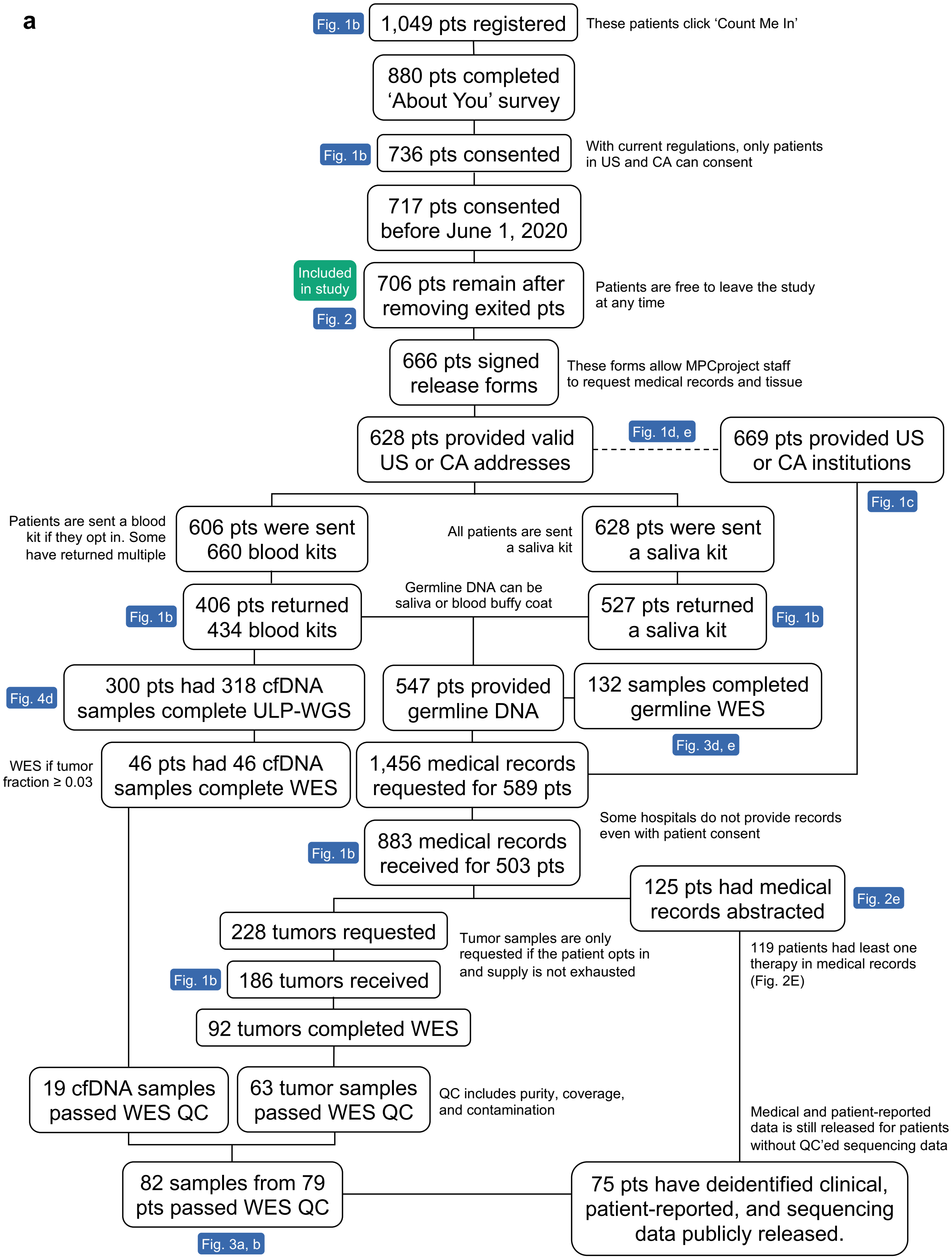
a
Fig. 1b
1,049 pts registered
These patients click 'Count Me In'
880 pts completed 'About You' survey
Fig. 1b
736 pts consented
With current regulations, only patients in US and CA can consent
717 pts consented before June 1, 2020
Included in study
Fig. 2
706 pts remain after removing exited pts
Patients are free to leave the study at any time
666 pts signed release forms
These forms allow MPCproject staff to request medical records and tissue
628 pts provided valid US or CA addresses
Fig. 1d, e
669 pts provided US or CA institutions
Fig. 1c
Patients are sent a blood kit if they opt in. Some have returned multiple
606 pts were sent 660 blood kits
All patients are sent a saliva kit
628 pts were sent a saliva kit
Fig. 1b
406 pts returned 434 blood kits
Germline DNA can be saliva or blood buffy coat
527 pts returned a saliva kit
Fig. 1b
Fig. 4d
300 pts had 318 cfDNA samples complete ULP-WGS
547 pts provided germline DNA
132 samples completed germline WES
Fig. 3d, e
WES if tumor fraction ≥ 0.03
46 pts had 46 cfDNA samples complete WES
1,456 medical records requested for 589 pts
Some hospitals do not provide records even with patient consent
Fig. 1b
883 medical records received for 503 pts
125 pts had medical records abstracted
Fig. 2e
228 tumors requested
Tumor samples are only requested if the patient opts in and supply is not exhausted
119 patients had least one therapy in medical records (Fig. 2E)
Fig. 1b
186 tumors received
92 tumors completed WES
19 cfDNA samples passed WES QC
63 tumor samples passed WES QC
QC includes purity, coverage, and contamination
Medical and patient-reported data is still released for patients without QC'ed sequencing data
82 samples from 79 pts passed WES QC
75 pts have deidentified clinical, patient-reported, and sequencing data publicly released.
Fig. 3a, b

**Supplementary Figure 6. MPCproject attrition chart**

**a)** Chart detailing project attrition for patient-partners that consented as of June 1, 2020. The chart represents data collected on June 7, 2021. Patient recruitment, sample acquisition, medical record abstraction, sequencing, and data releases are ongoing processes, so these values will grow as the project continues. Colored boxes indicate the figures that use those values in analysis and visualization. Values for Fig. 1b shown in this attrition chart may be greater than those shown in Fig. 1b at the study cutoff date, as Fig. 1b is a snapshot showing values collected as of June 1, 2020, while this attrition chart includes steps that may have been completed by consented patient-partners after June 1, 2020.

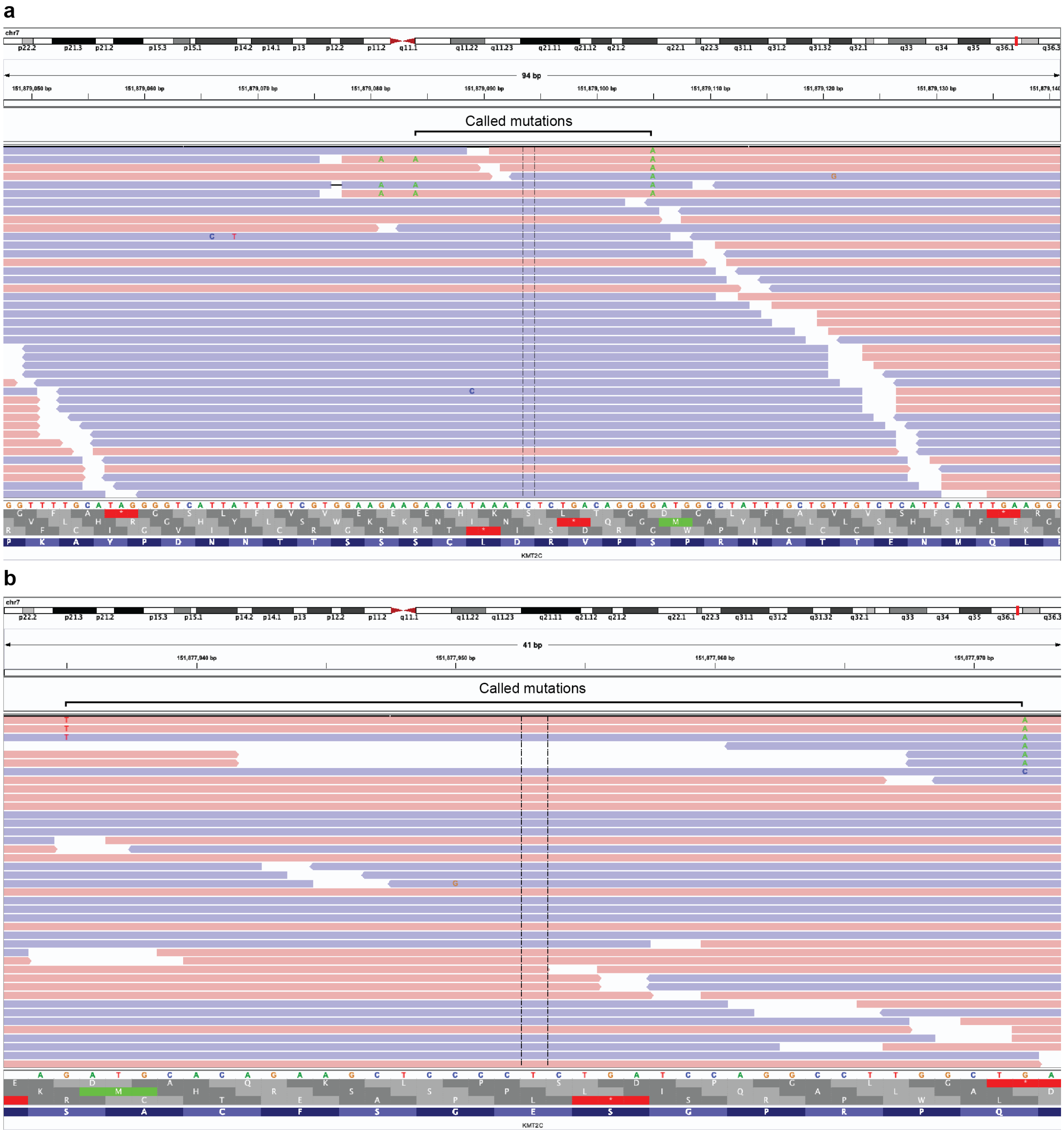


**Supplementary Figure 7. IGV screenshots of *KMT2C* mutation-sharing reads**

**a)** IGV screenshot containing reads that span somatic *KMT2C* mutations (chr7:151879084/p.S1947F and chr7:151879105/p.S1954F) in the cfDNA sample of patient-partner 0203. A mutation may also be present at chr7:151879081 but was rejected by Mutect’s internal filters as it is close to an inferred gap event. Coloring of reads indicates strand.

**b)** IGV screenshot containing reads that span somatic *KMT2C* mutations (chr7:151877972/ p.Q2325* and chr7:151877935/p.S2337Y). Coloring of reads indicates strand.

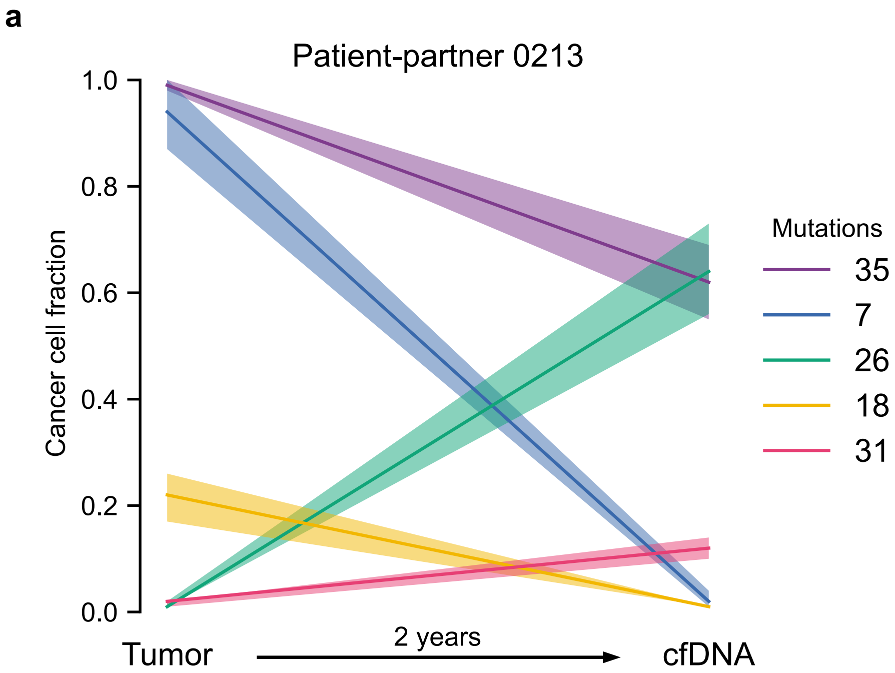


**Supplementary Figure 8. Phylogenetics of samples from patient-partner 0213**

**a)** The y-axis shows the cancer cell fraction (CCF) of clonal clusters identified between primary tumor and cfDNA from donated blood (x-axis). Colors indicate how many mutations were identified in each clone, with a 95% confidence interval around the estimated CCF. Purple represents the truncal/ancestral clone. The ancestral clone does not reach a CCF of 1 in the liquid sample because its inferred purity (0.20) is low, which confounds the ability to accurately quantify CCF.

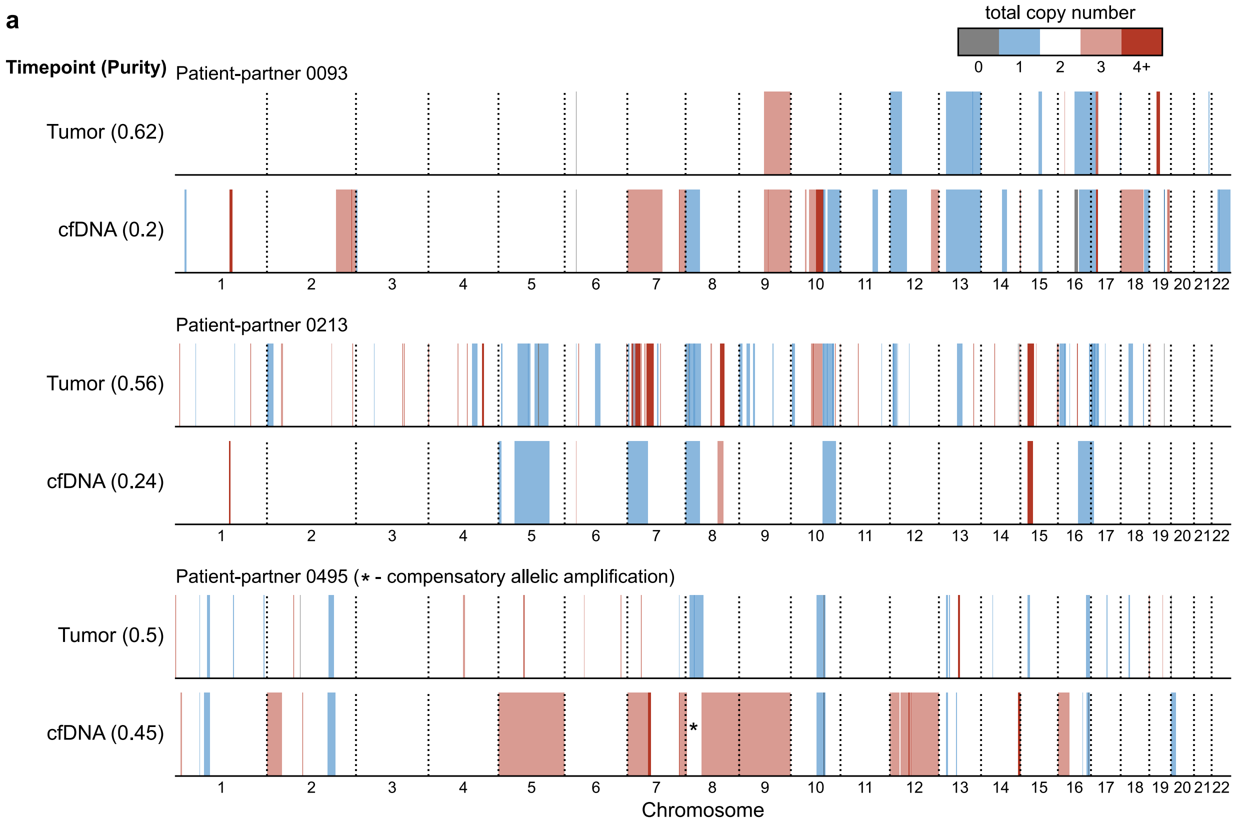


**Supplementary Figure 9. Copy number profiles of shared tumor and cfDNA samples**

**a)** Concordance of copy number profiles between archival primary tumors and donated cfDNA samples. The x-axis depicts chromosomal location, with coloring representing copy number alterations and their absolute copy number. In general, there are no archival-specific copy number alterations, with the potential exception of chr7p amplification in patient-partner 0213. When sample purity is below 0.30, focal copy number amplifications can be undetectable. In patient-partner 0495's samples, an arm-level deletion of 8p acquired a compensatory amplification on the other allele that restored diploid copy number.

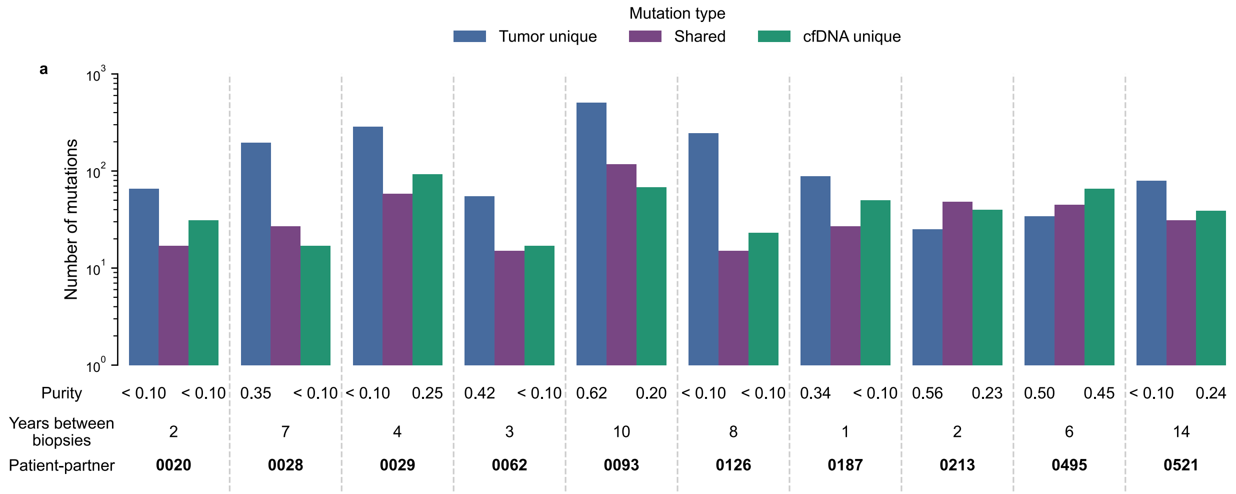


**Supplementary Figure 10. Mutation exclusivity between tumor and cfDNA samples from the same patient**

**a)** Number of mutations for each sample type for ten patient-partners with both archival tumor and donated cfDNA samples. The y-axis shows number of mutations, while the x-axis shows each patient. The purple and blue bars represent mutations identified exclusively in the archival tumor and cfDNA samples, respectively. The green bars represent mutations that had at least one supporting read in both tumor and cfDNA samples within the union of all mutations called in tumor and cfDNA samples (see Methods – *Phylogenetic analysis*). The purities and amount of time between samples are shown below each bar. Purities below 0.10 cannot be accurately estimated.

| Institution | Patient count | Institution | Patient count |
| --- | --- | --- | --- |
| DANA-FARBER CANCER INSTITUTE | 47 | UC HEALTH - UNIVERSITY OF COLORADO CANCER CENTER | 4 |
| UT M. D. ANDERSON CANCER CENTER | 29 | MASSEY | 3 |
| HELEN DILLER FAMILY COMPREHENSIVE CANCER CENTER | 26 | CARBONE | 3 |
| MAYO CLINIC HOSPITAL ROCHESTER | 24 | NORRIS COTTON | 3 |
| SIDNEY KIMMEL CANCER CENTER | 24 | COLUMBIA UNIVERSITY IRVING MEDICAL CENTER | 3 |
| MEMORIAL SLOAN | 19 | PERLMUTTER CANCER CENTER | 3 |
| HUTCHINSON | 17 | ROGEL | 3 |
| MOUNT SINAI HOSPITAL | 13 | STEPHENSON CANCER CENTER | 2 |
| SMILOW CANCER | 12 | ROSWELL PARK | 2 |
| KNIGHT | 11 | FOX CHASE | 2 |
| SITEMAN | 10 | CITY OF HOPE | 2 |
| MOORES CANCER CENTER | 10 | MASONIC | 2 |
| INGRAM CANCER | 8 | HOLLING | 2 |
| SIMON COMPREHENSIVE CANCER | 8 | SYLVESTER | 2 |
| NORTHWESTERN | 8 | HOLDEN | 2 |
| DUKE CANCER | 7 | OHIO STATE UNIVERSITY COMPREHENSIVE CANCER CENTER - THE JAMES | 1 |
| THE UNIVERSITY OF CHICAGO COMPREHENSIVE CANCER CENTER | 6 | HILLMAN | 1 |
| MOFFITT | 5 | LINEBERGER | 1 |
| UC DAVIS HEALTH - COMPREHENSIVE CANCER CENTER | 4 | CHAO FAMILY COMPREHENSIVE CANCER CENTER | 1 |
| UNIVERSITY OF KANSAS CANCER CENTER | 4 | UNIVERSITY OF NEW MEXICO | 1 |
| STANFORD CANCER INSTITUTE | 4 | BAYLOR | 1 |
| RUTGERS CANCER | 4 | MAYS | 1 |
| SIMMONS COMPREHENSIVE CANCER CENTER | 4 | UK MARKEY CANCER CENTER | 1 |
| WINSHIP | 4 | THOMAS JEFFERSON | 1 |
| KECK HOSPITAL OF USC - NORRIS CANCER CENTER | 4 | LOMBARDI | 1 |

**Supplementary Table 1. List of NCI-designated cancer centers**

List of NCI-designated cancer centers along with unique patient-partner attendance counts. For institutions that have satellite locations, only the main location was considered in tabulating patient attendance and NCI-designated status. These institutions are depicted in green in Fig. 1c.

| Patient-reported data | Number of patient-partners (%) |
|---|---|
| *Age at initial diagnosis (mean: 61)* | |
| Did not respond | 1 (0.1%) |
| ≤ 40 years | 4 (0.6%) |
| > 40, ≤ 50 years | 62 (8.7%) |
| > 50, ≤ 60 years | 256 (35.8%) |
| ≥ 60 years | 383 (54.8%) |
| *What is your race? (Select all that apply)* | |
| White | 657 (93.1%) |
| Black or African American | 12 (1.7%) |
| Other (Not specified) | 10 (1.4%) |
| Japanese | 4 (0.6%) |
| Chinese | 4 (0.6%) |
| American Indian | 3 (0.4%) |
| Prefer to not respond | 3 (0.4%) |
| Did not respond | 4 (0.4%) |
| Southeast Asian or Indian | 2 (0.3%) |
| American Indian and White | 2 (0.3%) |
| White, Other (Not specified) | 2 (0.3%) |
| Japanese and White | 2 (0.3%) |
| Japanese, Chinese, Hawaiian, and White | 1 (0.1%) |
| *Do you consider yourself Hispanic or Latino?* | |
| Yes | 12 (1.7%) |
| No | 689 (97.6%) |
| Did not respond | 5 (0.7%) |

**Supplementary Table 2. Additional patient reported data**

Patient reported demographic data for patient-partners enrolled before June 1, 2020 (n = 706). Age at initial prostate cancer diagnosis is calculated based on the patient reported date of birth and month/year of initial prostate cancer diagnosis. Patient-partners were free to select as many racial identities as they identified with.

| Gene | DNA-repair | Germline susceptibility |
|---|---|---|
| *ATM* | X | X |
| *ATR* | X | |
| *BAP1* | X | |
| *BARD1* | X | |
| *BRCA1* | X | X |
| *BRCA2* | X | X |
| *BRIP1* | X | |
| *CDK12* | X | |
| *CHEK1* | X | |
| *CHEK2* | X | X |
| *ERCC3* | X | |
| *FAM175A* | X | |
| *FANCA* | X | |
| *FANCL* | X | |
| *FANCM* | X | |
| *GEN1* | X | |
| *HDAC2* | X | |
| *HOXB13* | | X |
| *MLH1* | X | X |
| *MLH3* | X | |
| *MRE11A* | X | |
| *MSH2* | X | X |
| *MSH6* | X | X |
| *NBN* | X | X |
| *PALB2* | X | |
| *PMS2* | X | |
| *PPP2R2A* | X | |
| *RAD50* | X | |
| *RAD51* | X | |
| *RAD51B* | X | |
| *RAD51C* | X | |
| *RAD51D* | X | |
| *RAD54L* | X | |
| *XRCC2* | X | |

**Supplementary Table 3. DNA repair and germline susceptibility gene list**

List of genes used in the analysis of the association between the presence of COSMIC2.0 signature 3 and DNA-repair alterations (DNA-repair) and list of genes used to evaluate germline alterations in prostate cancer susceptibility genes. An X signifies that the gene was used in that analysis. DNA-repair related genes taken from Mateo et al. 2015, de Bono et al. 2020, and Pritchard et al. 2016[1–3]. Prostate cancer susceptibility genes taken from Aldubayan 2019[4]. See Methods for the specifics of these analyses.

| Therapy brand name (Generic name) | Category | Number of patient-partners (% of 639) |
|---|---|---|
| *Hormones* | | |
| Lupron (Leuprolide) | 1st line ADT | 538 (84.2%) |
| Zoladex (Goserelin) | 1st line ADT | 38 (5.9%) |
| Casodex (Bicalutamide) | 1st line ADT | 326 (51.0%) |
| Drogenil (Flutamide) | 1st line ADT | 5 (0.8%) |
| Nilandron (Nilutamide) | 1st line ADT | 5 (0.8%) |
| Xtandi (Enzalutamide) | 2nd line ADT | 107 (16.7%) |
| Zytiga (Abiraterone) | 2nd line ADT | 220 (34.4%) |
| Prostap (Leuprorelin) | 1st line ADT | 1 (0.2%) |
| Firmagon (Degarelix) | 1st line ADT | 109 (17.1%) |
| Suprefact (Buserelin) | 1st line ADT | 0 (0.0%) |
| Decapeptyl (Triptorelin) | 1st line ADT | 3 (0.4%) |
| *Chemotherapy* | | |
| Taxotere (Docetaxel) | Chemotherapy | 168 (26.3%) |
| Taxol (Paclitaxel) | Chemotherapy | 2 (0.3%) |
| Paraplatin (Carboplatin) | Chemotherapy | 17 (2.6%) |
| Etopophos / Toposar (Etoposide) | Chemotherapy | 5 (0.8%) |
| Novantrone (Mitoxantrone) | Chemotherapy | 1 (0.2%) |
| Emcyt (Estramustine) | Chemotherapy | 3 (0.5%) |
| Jevtana (Cabazitaxel) | Chemotherapy | 18 (2.8%) |
| *Other Therapy* | | |
| Provenge (Sipuleucel-T) | Immunotherapy | 59 (9.2%) |
| Opdivo (Nivolumab) | Immunotherapy | 2 (0.3%) |
| Keytruda (Pembrolizumab) | Immunotherapy | 10 (1.5%) |
| Yervoy (Ipilimumab) | Immunotherapy | 3 (0.5%) |
| Tecentriq (Atezolizumab) | Immunotherapy | 0 (0.0%) |
| Lynparza (Olaparib) | PARP inhibitor | 6 (0.9%) |
| Rubraca (Rucaparib) | PARP inhibitor | 0 (0.0%) |
| Xofigo (Radium-223) | Nuclear medicine | 23 (3.6%) |
| Zometa (Zoledronic Acid) | Supportive care | 50 (7.9%) |
| Xgeva/Prolia (Denosumab) | Supportive care | 103 (16.2%) |
| Quadramet (Samarium SM 153 lexidronam) | Supportive care | 0 (0.0%) |
| Metastron (Strontium-89) | Supportive care | 0 (0.0%) |
| *Experimental/Clinical Trial* | | |
| Experimental/Clinical Trial | Clinical trial | 87 (13.6%) |

**Supplementary Table 4. Therapies available for selection on patient survey**

List of therapies available for selection on patient survey (Supplementary Figure 2). Only these therapies were used to determine the overlap between patient-reported therapies and medical record therapies. Percentage defined relative to the number of patient-partners that provided at least one therapy on the survey (n = 639/706).
