## Supplementary Methods for "A patient-driven clinicogenomic partnership through the Metastatic Prostate Cancer Project"

### Table of Contents

**Patient Enrollment and Study Material Acquisition** 2
- Establishing patient partnership 2
- Patient Enrollment and Informed Consent 2
- Medical Records 3
- Samples 3
  - Saliva 3
  - Archived Tumor Tissue 3
  - Primary and Secondary Blood Samples 4

**Data Generation** 4
- Medical Record Abstraction 4
- Patient-Reported Data 5
  - Study inclusion 5
  - Cleaning/categorization 5
    - Medical Institutions 5
    - Therapies 5
    - Alternative lifestyles 4
- Genomic Sequencing 5
  - DNA Isolation 5
    - Saliva 5
    - cfDNA Extraction from Whole Blood 6
  - Ultra-Low Pass Whole Genome Sequencing (ULP-WGS) 6
    - non-UMI ULP-WGS sequencing [dates: 2017-2/11/2018] 6
    - UMI ULP-WGS sequencing [dates: 2/12/2018-6/1/2020] 7
  - Whole Exome Sequencing (WES) 7
    - Express WES for saliva and tissue [dates: 8/13/2017 - 4/15/2018] 7
    - Express WES for saliva and tissue [dates: 4/15/2018-6/1/2020] 8
    - Deep ICE Exome from Non-UMI Enabled ULP Libraries Methods [dates: previous to 8/13/2017] 9
    - Deep ICE Exome from UMI-Enabled ULP Libraries [dates: 8/13/2017-6/1/2020] 9

**References** 10

### *Patient Enrollment and Study Material Acquisition*

#### Establishing patient partnership

Patients and the extended metastatic prostate community have been directly involved in the creation and development of the Metastatic Prostate Cancer Project (MPCproject) since the project's conception. During the initial development of the project, a patient advisory council (PAC) comprised of patients, loved ones, and advocates met frequently with study staff to determine the study's approach for outreach, patient enrollment, study website design, and sample collection, among other details of project operations. Study staff from the project continue to meet regularly with the PAC. In addition to working with members of the PAC, the MPCproject leverages the expertise of the many prostate cancer advocacy group partners to improve outreach and project operations. Finally, patients that are not directly involved in the PAC or an advocacy group, can learn about and partner with the project through various social media platforms, newsletters, or educational materials generated by study staff to provide input or feedback.

This study includes as authors patient advocates who were instrumental in survey design, project development, assessment of patient criteria, and outreach strategy. The MPCproject glossary included with the study was reviewed by practicing oncologists, patient advocates, and study staff.

#### Patient Enrollment and Informed Consent

The MPCproject is a decentralized, online patient-partnered genomics research study. Patients anywhere in the United States and Canada can visit the project webiste (https://mpcproject.org) to learn about the research initiative and register for the study. If a patient is interested in participating, the online registration process has four steps: registration, an optional intake survey, an electronic consent form, and a medical record release form. For the study, we consider any patient that completes the consent form to be enrolled.

On the study registration page, a patient provides their first and last name, email, and confirmation of their metastatic or advanced prostate cancer diagnosis as well as acknowledgement of their willingness to provide further information on their medical care and experience with the disease. The registration page prompts patients to create a password protected account to save provided information and to allow patients to revisit their completed survey and forms at any time. Once the account has been created, registrants are taken to an optional intake survey (Supplemental Figure 2) where they are asked to provide basic demographic information as well as answer questions about their experience with prostate cancer via a 17-question survey that was developed in partnership with clinicians, researchers, and patients. Each question is optional and survey responses can be revisited. To submit the survey, patients agree to the MPCproject saving their survey information, and, if they live in the U.S. or Canada, agree to study staff reaching out if the MPCproject conducts future studies. The minimum requirement to submit the survey is providing country of origin and a zip code.

Registrants that choose to submit the survey and who reside in the U.S. or Canada are then taken to an electronic consent form. Patients provide informed consent using a web-based consent form as approved by the Dana-Farber/Harvard Cancer Center Institutional Review Board (DF/HCC Protocol 15-057B). To formally enroll in the study, patients provide their electronic signature on the consent form. The consent form provides various levels of participation. The minimum consent enables study staff to request and abstract medical records, send the patient a saliva kit, perform germline sequencing analysis if a saliva sample is returned, and release de-identified clinical and genomic data into public repositories. Patients have the additional option of consenting to study staff obtaining archived tumor tissue and/or blood sample(s) for further somatic and germline sequencing analyses. Email reminders are sent to registrants who have not completed the consent process (weekly for three weeks, and again at six weeks). A copy of the completed consent form is saved in the patient's account and emailed to them.

Upon submission of the consent form, the final step in the study enrollment process is to complete a medical release form. On this form, patients provide their contact information and information about any physician or hospital involved in the care of their prostate cancer. By submitting the release form, patients agree to study staff reaching out to the listed institutions to requested medical records and, if elected on the electronic consent form, archived tissue samples. Email reminders are sent weekly for three weeks, and again at six weeks, to registrants who have not completed the release form. A copy of the completed release form is saved in the patient's account and emailed to them.

### Medical Records

After patients complete the consent and release forms and provide institutions where they received care for their prostate cancer, the study staff requests their medical records. Study staff call each institution's medical record departments to obtain copies of the patient's records starting at the date of diagnosis of prostate cancer through the day of the faxed request. Requests are faxed to the respective departments after phone confirmation of the fax number. Medical records are returned to the project via mail, fax, or online portals. Once a medical record arrives, it is saved in an electronic format in a secure database. If a record request is not fulfilled in 6 months, a second request is submitted. If the medical records department requires additional paperwork or signatures per the specific institution's release requirements, the patient is contacted and asked to provide the additional required forms. When patients are contacted for this purpose, study staff are clear that this additional step is optional for patients. Study staff can also request subsequent medical records after an initial request had been fulfilled if the need arises.

### Samples

All patients that complete the electronic consent form are sent a saliva kit to provide a saliva sample. In addition, patients can opt-in to providing archival tumor tissue and/or one or more blood samples.

#### *Saliva*

Saliva kits are sent to patients who complete the consent and medical release form and provide a valid mailing address in the United States or Canada. Staff at the Broad Institute Genomics Platform prepare each unique patient's kit by assigning it a unique barcode and prepaid business reply-label and packaging the kit with instructions for the patient on how to provide at least 2 mL of saliva in a DNA Genotek Oragene Discover (OGR-600) tube labeled with a matching barcode. All kits are affixed with a prepaid business-reply label. Samples are mailed back to the Broad Institute by patients after collection, and then logged and stored at room temperature by study staff upon receipt. Saliva samples are eventually pushed for whole exome sequencing to obtain germline DNA once matched tumor samples are also received and submitted for sequencing.

#### *Archived Tumor Tissue*

Once a patient's medical record and normal normal sample (saliva or blood) are received, study staff review the record to confirm the patient has had a clinical diagnosis of metastatic or advanced prostate cancer. Surgical and pathology records are used to develop a patient's surgical history and identify archived formalin-fixed paraffin embedded (FFPE) prostate cancer tumor tissue that may be requested. Study staff, in collaboration with oncologists and pathologists, developed strict guidelines for selecting which tumor sample to request to obtain the minimal amount of tissue that will not interfere with the patient's future clinical care. For each patient, a specific sample is requested only there are at least three blocks with prostatic adenocarcinoma and at minimum two of those blocks are actively being stored in the source pathology department. If a sample meets the requesting criteria, study staff coordinate with the sending pathology department to fax a request and obtain the sample via mail. The tissue request form requests that pathology departments send an H&E slide along with either an entire block from the surgical case or 5-20 5-micron unstained slides from a block. All tissue requests submitted by the MPCproject state that no sample should be exhausted to fulfill the request. Tissue samples received as blocks are labeled with unique numerical identifiers and sent to the Dana-Farber/Harvard Cancer Center Specialized Histopathology Services (SHS) Core to be cut into three 30- micron scrolls per block and an accompanying H&E for tumor confirmation. Scrolls, unstained slides, and H&Es are labeled with unique barcode identifiers. Archived tumor tissue with a matched germline sample (from either saliva or a blood sample's buffy coat) are sent to the Broad Institute's Genomics Platform for whole exome sequencing.

*Primary and Secondary Blood Samples*
Blood sample acquisition and sequencing preparation are performed as described in Painter et al. 2020 except in the additional steps of sending secondary blood kits to patients[1]. The MPCproject was awarded a grant to send a cohort of selected patients second blood kits to obtain an additional blood sample to study tumor evolution. Patients are selected based on a combination of criteria including date of registration, date of primary blood draw, primary blood sample containing sufficient ctDNA quantity for whole exome sequencing, and successful acquisition of medical records. An email is sent to selected patients describing the intent and optional nature of the second blood kit. The email contains a link to a new consent form and asks if they would be willing to provide an additional blood sample. If the patient selects 'Yes' on the consent form, another round of the blood sample acquisition process is triggered: a new blood kit is sent to their home, returned to the Broad Institution, and processed using the same procedure outlined for their primary kit.

### ***Data Generation***

#### Medical Record Abstraction

Medical records are requested for any consented patient in the US and Canada that listed any institution(s) from which they received care on their medical release form. Medical records arrive in various formats and all are eventually transferred to an electronic format and stored on a secure internal server. Scanned medical records are run through the Optical Character Recognition (OCR) engine known as Tesseract (LSTM model inside Tesseract version 4.0; (https://github.com/tesseract-ocr/tesseract)) to facilitate manual abstraction by study staff.

Three separate abstractors on the study staff team are involved in the abstraction and quality control process of the clinical data from each searchable record. To begin, two abstractors independently read and isolate the same clinical information for each patient. A third abstractor aligns the separate abstractions and identifies field-specific discrepancies between the two abstractions. The third abstractor attempts to resolve any lack of concordance by returning to the patient's medical record to identify the correct data. At any point in the process, abstractors can work with clinical oncologists to answer questions or address lack of concordance.

The abstractors use a clinical data dictionary comprising 60 fields that was curated by prostate cancer oncologists. For information that's not found, it was abstracted as 'NOT FOUND IN RECORD'. In instances where ambiguity or incomplete data was present, inferences were made considering the whole narrative of the medical record. The dictionary includes possible responses for each field. For date-type fields, incomplete dates missing either the month or day, are abstracted as the first month of the year and/or first day of the month, respectively. All time related fields are anchored from the date of primary prostate cancer diagnoses. For example, a patient's metastatic diagnosis date is represented as the calculated number of days from the primary diagnosis date to the metastatic diagnosis date. This was done to protect patient privacy.

### Patient-Reported Data

#### *Study inclusion*

Survey responses were cleaned for patients that completed their consent and release forms and submitted a survey by June 1, 2020. 706 of these patients reported being located within the U.S. and Canada and were thus included in downstream analyses.

#### *Cleaning/categorization*

**Medical Institutions**

Patients were asked in their medical release form to report all physicians with whom they received care for their prostate cancer, institutions where they received an initial prostatic biopsy or prostatectomy, and institutions where they received another surgery related to their prostate cancer. Institutions of reported physicians were gathered based on the most recent affiliation identified from affiliated websites. Satellite locations of larger institutions were considered separate institutions. Names were standardized by three separate reviewers manually. For the purpose of Fig 1c, only unique institutions for each patient are shown. The NCI designated cancer center list was taken from cancer.gov/research/infrastructure/cancer-centers/find.

**Therapies**

Patients selected all therapies that they had received for their prostate cancer in the intake survey. Therapies were categorized by prostate cancer oncologists into broad treatment categories according to their primary therapeutic function (See Supplementary Table 4).

**Alternative lifestyles**

Patient responses to question 7 on the intake survey (Supplementary Fig 2) were categorized into four broad categories: Diet/lifestyle, Supplements, and Non-Cancer Therapies. Except for plant-based diet and unspecified diet change, responses were not mutually exclusive. Different methods of taking similar supplements (e.g., turmeric paste, turmeric capsules, turmeric powder) were considered the same supplement. Brand name products were converted to generic forms (e.g., Pomi-T was considered "pomegranate"). Manual classification was conducted by two separate reviewers.

### Genomic Sequencing

All samples were received and sequenced at the Broad Institute's Genomics Platform. Due to changes in sequencing methods as a function of improved technologies and the longitudinal nature of this project, certain sequencing methods are subset by date to indicate what was applied for samples received within the specific timeframe.

#### *DNA Isolation*

**Saliva**

DNA was extracted via the Chemagic MSM I with the Chemagic DNA Blood Kit-96 from Perkin Elmer. This kit combines a chemical and mechanical lysis with magnetic bead-based purification. Saliva samples were incubated at 50°C for 2 hours. The saliva was then transferred to a deep well plate placed on the Chemagic MSM I. The following steps were automated on the MSM I.

M-PVA Magnetic Beads were added to the saliva. Lysis buffer was added to the solution and mixed. The bead-bound DNA was then removed from solution via a 96-rod magnetic head and washed in three Ethanol-based wash buffers. The beads were then washed in a final water wash buffer. Finally, the beads were dipped in elution buffer to resuspend the DNA sample in solution. The beads were then removed from solution, leaving purified DNA eluate. DNA samples were quantified using a fluorescence based PicoGreen assay.

**cfDNA Extraction from Whole Blood**

Whole blood was collected in EDTA, CellSave, or Streck tubes and processed for plasma fractionation. Blood tubes were centrifuged at 1900 g for 10 minutes and plasma was transferred to second tube before further centrifugation at 15000 g for 10 minutes. Supernatant plasma was stored at -80C until cfDNA extraction. cfDNA was extracted using the QIAsymphony DSP Circulating DNA Kit according to the manufacturer's instructions, with 6.3 mL of plasma as input and with a 60 uL DNA elution (Qiagen, 2017).

*Ultra-Low Pass Whole Genome Sequencing (ULP-WGS)*

**non-UMI ULP-WGS sequencing [dates: 2017-2/11/2018]**

1. Library Construction
Initial DNA input is normalized to be within the range of 25-52.5 ng in 50 uL of TE buffer (10mM Tris HCl 1mM EDTA, pH 8.0) according to picogreen quantification. For adapter ligation, Illumina paired end adapters were replaced with palindromic forked adapters, purchased from Integrated DNA Technologies, with unique dual-indexed molecular barcode sequences to facilitate downstream pooling. With the exception of the palindromic forked adapters, the reagents used for end repair, A-base addition, adapter ligation, and library enrichment PCR were purchased from KAPA Biosciences in 96-reaction kits. In addition, during the post-enrichment SPRI cleanup, elution volume was reduced to 30µL to maximize library concentration, and a vortexing step was added to maximize the amount of template eluted.

2. Post Library Construction Quantification and Normalization
Library quantification was performed using the Invitrogen Quant-It broad range dsDNA quantification assay kit (Thermo Scientific Catalog: Q33130) with a 1:200 PicoGreen dilution. Following quantification, each library is normalized to a concentration of 25 ng/µL, using a 1X Low TE pH 7.0 solution.

3. Library Pool Creation for Ultra-low Pass Sequencing
In preparation for the sequencing of the ultra-low pass libraries (ULP), approximately 4 µL of the normalized library is transferred into a new receptacle and further normalized to a concentration of 2ng/µL using Tris-HCl, 10mM, pH 8.0. Following normalization, up to 95 ultra-low pass WGS samples are pooled together using equivolume pooling. The pool is quantified via qPCR and normalized to the appropriate concentration to proceed to sequencing.

4. Cluster amplification and sequencing
Cluster amplification of library pools was performed according to the manufacturer's protocol (Illumina) using Exclusion Amplification cluster chemistry and HiSeq X flowcells. Flowcells were sequenced on v2 Sequencing-by-Synthesis chemistry for HiSeq X flowcells. The flowcells are then analyzed using RTA v.2.7.3 or later. Each pool of ultra-low pass whole genome libraries is run on one lane using paired 151bp runs.

**UMI ULP-WGS sequencing [dates: 2/12/2018-6/1/2020]**

1. Library Construction
Initial DNA input is normalized to be within the range of 25-52.5 ng in 50 uL of TE buffer (10mM Tris HCl 1mM EDTA, pH 8.0) according to picogreen quantification. Library preparation is performed using a commercially available kit provided by KAPA Biosystems (KAPA HyperPrep Kit with Library Amplification product KK8504) and IDT's duplex UMI adapters. Unique 8-base dual index sequences embedded within the p5 and p7 primers (purchased from IDT) are added during PCR. Enzymatic clean-ups are performed using Beckman Coultier AMPure XP beads with elution volumes reduced to 30µL to maximize library concentration.

2. Post Library Construction Quantification and Normalization
Library quantification was performed using the Invitrogen Quant-It broad range dsDNA quantification assay kit (Thermo Scientific Catalog: Q33130) with a 1:200 PicoGreen dilution. Following quantification, each library is normalized to a concentration of 35 ng/µL, using Tris-HCl, 10mM, pH 8.0.

3. Library Pool Creation for Ultra-low Pass Sequencing
In preparation for the sequencing of the ultra-low pass libraries (ULP), approximately 4 µL of the normalized library is transferred into a new receptacle and further normalized to a concentration of 2ng/µL using Tris-HCl, 10mM, pH 8.0. Following normalization, up to 95 ultra-low pass WGS samples are pooled together using equivolume pooling. The pool is quantified via qPCR and normalized to the appropriate concentration to proceed to sequencing.

4. Cluster amplification and sequencing
Cluster amplification of library pools was performed according to the manufacturer's protocol (Illumina) using Exclusion Amplification cluster chemistry and HiSeq X flowcells. Flowcells were sequenced on v2 Sequencing-by-Synthesis chemistry for HiSeq X flowcells. The flowcells are then analyzed using RTA v.2.7.3 or later. Each pool of ultra-low pass whole genome libraries is run on one lane using paired 151bp runs

### *Whole Exome Sequencing (WES)*

**Express WES for saliva and tissue [dates: 8/13/2017 - 4/15/2018]**

1. Library Construction
Library construction was performed as described in Fisher et al., with the following modifications DNA input into shearing was reduced from 3µg to 10-100ng in 50µL of solution. For adapter ligation, Illumina paired end adapters were replaced with palindromic forked adapters, purchased from Integrated DNA Technologies, with unique dual-indexed molecular barcode sequences to facilitate downstream pooling. Kapa HyperPrep reagents in 96-reaction kit format were used for end repair/A-tailing, adapter ligation, and library enrichment PCR. In addition, during the post-enrichment SPRI cleanup, elution volume was reduced to 30µL to maximize library concentration, and a vortexing step was added to maximize the amount of template eluted.

2. In-solution hybrid selection
After library construction, hybridization and capture were performed using the relevant components of Illumina's TruSeq Rapid Exome Kit and following the manufacturer's suggested protocol, with the following exceptions: first, all libraries within a library construction plate were pooled prior to hybridization. Second, the Midi plate from Illumina's TruSeq Rapid Exome Kit was replaced with a skirted PCR plate to facilitate automation. All hybridization and capture steps were automated on the Agilent Bravo liquid handling system.

3. Preparation of libraries for cluster amplification and sequencing
After post-capture enrichment, library pools were quantified using qPCR (automated assay on the Agilent Bravo), using a kit purchased from KAPA Biosystems with probes specific to the ends of the adapters. Based on qPCR quantification, libraries were normalized to 2nM, then denatured using 0.1 N NaOH on the Hamilton Starlet. After denaturation, libraries were diluted to 20pM using hybridization buffer purchased from Illumina.

4. Cluster amplification and sequencing
Cluster amplification of denatured templates was performed according to the manufacturer's protocol (Illumina) using HiSeq 4000 cluster chemistry and HiSeq 4000 flowcells. Flowcells were sequenced on v1 Sequencing-by-Synthesis chemistry for HiSeq 4000 flowcells. The flowcells are then analyzed using RTA v.1.18.64 or later. Each pool of whole exome libraries was run on paired 76bp runs, reading the dual-indexed sequences to identify molecular indices and sequenced across the number of lanes needed to meet coverage for all libraries in the pool.

**Express WES for saliva and tissue [dates: 4/15/2018-6/1/2020]**

1. Library Construction
Library construction was performed as described in Fisher et al., with the following modifications: initial genomic DNA input into shearing was reduced from 3μg to 10-100ng in 50μL of solution. For adapter ligation, Illumina paired end adapters were replaced with palindromic forked adapters, purchased from Integrated DNA Technologies, with unique dual-indexed molecular barcode sequences to facilitate downstream pooling. Kapa HyperPrep reagents in 96-reaction kit format were used for end repair/A-tailing, adapter ligation, and library enrichment PCR. In addition, during the post-enrichment SPRI cleanup, elution volume was reduced to 30μL to maximize library concentration, and a vortexing step was added to maximize the amount of template eluted.

2. In-solution hybrid selection
After library construction, hybridization and capture were performed using the relevant components of Illumina's TruSeq Rapid Exome Kit and following the manufacturer's suggested protocol, with the following exceptions: first, all libraries within a library construction plate were pooled prior to hybridization. Second, the Midi plate from Illumina's TruSeq Rapid Exome Kit was replaced with a skirted PCR plate to facilitate automation. All hybridization and capture steps were automated on the Agilent Bravo liquid handling system.

3. Preparation of libraries for cluster amplification and sequencing
After post-capture enrichment, library pools were quantified using qPCR (automated assay on the Agilent Bravo), using a kit purchased from KAPA Biosystems with probes specific to the ends of the adapters. Based on qPCR quantification, libraries were normalized to 2nM, then denatured using 0.2 N NaOH on the Hamilton Starlet. After denaturation, libraries were diluted to 20pM using hybridization buffer purchased from Illumina.

4. Cluster amplification and sequencing
Cluster amplification of denatured templates was performed according to the manufacturer's protocol (Illumina) using exclusion amplification cluster chemistry and HiSeq X flowcells. Flowcells were sequenced on v2.5 Sequencing-by-Synthesis chemistry for HiSeq X flowcells. The flowcells are then analyzed using RTA v.2.7.0 or later. Each pool of whole exome libraries was run on paired 76bp runs, reading the dual-indexed sequences to identify molecular indices and sequenced across the number of lanes needed to meet coverage for all libraries in the pool.

**Deep ICE Exome from Non-UMI Enabled ULP Libraries Methods [dates: previous to 8/13/2017]**

1. Library Construction
Initial DNA input is normalized to be within the range of 25-52.5 ng in 50 uL of TE buffer (10mM Tris HCl 1mM EDTA, pH 8.0) according to picogreen quantification. For adapter ligation, Illumina paired end adapters were replaced with palindromic forked adapters, purchased from Integrated DNA Technologies, with unique dual-indexed molecular barcode sequences to facilitate downstream pooling. With the exception of the palindromic forked adapters, the reagents used for end repair, A-base addition, adapter ligation, and library enrichment PCR were purchased from KAPA Biosciences in 96-reaction kits. In addition, during the post-enrichment SPRI cleanup, elution volume was reduced to 30μL to maximize library concentration, and a vortexing step was added to maximize the amount of template eluted.

2. In-solution hybrid selection
After library construction, hybridization and capture were performed using the relevant components of Illumina's Nextera Rapid Capture Exome Kit and following the manufacturer's suggested protocol, with the following exceptions: first, all libraries within a library construction plate were pooled prior to hybridization. Second, the Midi plate from Illumina's Nextera Rapid Capture Exome Kit was replaced with a skirted PCR plate to facilitate automation. All hybridization and capture steps were automated on the Agilent Bravo liquid handling system.

3. Preparation of libraries for cluster amplification and sequencing
After post-capture enrichment, library pools are quantified using qPCR (automated assay on the Agilent Bravo), using a kit purchased from KAPA Biosystems with probes specific to the ends of the adapters. Based on qPCR quantification, pools are normalized using a Hamilton Starlet to 2nM and sequenced using Illumina sequencing technology.

4. Cluster amplification and sequencing
Cluster amplification of library pools was performed according to the manufacturer's protocol (Illumina) using Exclusion Amplification cluster chemistry and HiSeq X flowcells. Flowcells were sequenced on v2 Sequencing-by-Synthesis chemistry for HiSeq X flowcells. The flowcells are then analyzed using RTA v.2.7.3 or later. Each pool of libraries was run on paired 151bp runs, reading the dual-indexed sequences to identify molecular indices and sequenced across the number of lanes needed to meet coverage for all libraries in the pool.

**Deep ICE Exome from UMI-Enabled ULP Libraries [dates: 8/13/2017-6/1/2020]**

1. Library Construction
Initial DNA input is normalized to be within the range of 25-52.5 ng in 50 uL of TE buffer (10mM Tris HCl 1mM EDTA, pH 8.0) according to picogreen quantification. Library preparation is performed using a commercially available kit provided by KAPA Biosystems (KAPA HyperPrep Kit with Library Amplification product KK8504) and IDT's duplex UMI adapters. Unique 8-base dual index sequences embedded within the p5 and p7 primers (purchased from IDT) are added during PCR. Enzymatic clean-ups are performed using Beckman Coultier AMPure XP beads with elution volumes reduced to 30μL to maximize library concentration.

2. Post Library Construction Quantification and Normalization
Library quantification was performed using the Invitrogen Quant-It broad range dsDNA quantification assay kit (Thermo Scientific Catalog: Q33130) with a 1:200 PicoGreen dilution. Following quantification, each library is normalized to a concentration of 25 ng/μL, using Tris-HCl, 10mM, pH 8.0.

3. In-solution hybrid selection

After library construction, hybridization and capture were performed using the relevant components of Illumina's TruSeq Rapid Exome Kit and following the manufacturer's suggested protocol, with the following exceptions: first, all libraries within a library construction plate were pooled prior to hybridization. Second, the Midi plate from Illumina's TruSeq Rapid Exome Kit was replaced with a skirted PCR plate to facilitate automation. All hybridization and capture steps were automated on the Agilent Bravo liquid handling system.

4. Preparation of libraries for cluster amplification and sequencing
After post-capture enrichment, library pools are quantified using qPCR (automated assay on the Agilent Bravo), using a kit purchased from KAPA Biosystems with probes specific to the ends of the adapters. Based on qPCR quantification, pools are normalized using a Hamilton Starlet to 2nM and sequenced using Illumina sequencing technology.

5. Cluster amplification and sequencing
Cluster amplification of library pools was performed according to the manufacturer's protocol (Illumina) using Exclusion Amplification cluster chemistry and HiSeq X flowcells. Flowcells were sequenced on v2 Sequencing-by-Synthesis chemistry for HiSeq X flowcells. The flowcells are then analyzed using RTA v.2.7.3 or later. Each pool of libraries was run on paired 151bp runs, reading the dual-indexed sequences to identify molecular indices and sequenced across the number of lanes needed to meet coverage for all libraries in the pool.

### ***References***


1. Painter, C. A. *et al.* The Angiosarcoma Project: enabling genomic and clinical discoveries in a rare cancer through patient-partnered research. *Nat. Med.* **26**, 181–187 (2020)

2. Fisher S. et al. A scalable, fully automated process for construction of sequence-ready human exome targeted capture libraries. Genome Biology 12, R1 (2011).
